# Analytical Comparability to Evaluate Impact of Manufacturing Changes of ARX788, an Anti-HER2 ADC in Late-Stage Clinical Development

**DOI:** 10.1101/2023.03.27.534450

**Authors:** Wayne Yu, Mysore Ramprasad, Manoj Pal, Chris Chen, Shashi Paruchuri, Lillian Skidmore, Nick Knudsen, Molly Allen, Ying Buechler

## Abstract

ARX788 is an anti-HER2 antibody drug conjugate (ADC) developed using Ambrx proprietary Engineered Precision Biologics technology. The manufacturing process of ARX788 has been optimized during the course of early to late-phase clinical development. A comprehensive evaluation of side-by-side comparability between pre- and post-change process for ARX788 drug substance and drug product from a quality perspective was conducted based on ICH Q5E guidelines consisting of batch release assays, physicochemical and biophysical characterization, biological characterization, and forced degradation studies. All results have demonstrated a high degree of similarity between the pre- and post-change ARX788 drug substance batches and drug product lots, demonstrating that the process manufacturing changes did not impact product quality.

## Introduction

Antibody-drug conjugates (ADCs) are a class of therapeutics that combines an antigen-specific antibody backbone with a potent cytotoxic payload, resulting in an improved therapeutic index. The development of anti-human epidermal growth factor receptor 2 (HER2) agents has been one of the most meaningful advancements in the management of metastatic breast cancer, significantly improving survival outcomes [1]. Two anti-HER2 ADCs have been approved by the FDA in HER2-positive breast cancer.

ARX788 is a next-generation, site-specific anti-HER2 ADC currently in global clinical trials for the treatment of HER2-positive metastatic breast cancer and gastric cancer [2–6]. It is comprised of a humanized HER2-targeting monoclonal antibody (mAb), conjugated to a cytotoxic tubulin inhibitor payload linker, Amberstatin (AS269), via a nonnatural amino acid incorporated into the antibody. The highly site-specific conjugation technology allows for a very stable oxime bond formed by conjugating a hydroxylamine containing drug-linker with the nonnatural amino acid, para-acetylphenylalanine (pAF), incorporated into the anti-HER2 mAb. This results in exactly two drug-linkers conjugated site specifically to one antibody molecule [2]. Further, the Ambrx Engineered Precision Biologics (EPB) platform technology with ARX788 offers unique advantages with superior efficacy, stability in systemic circulation, and a favorable safety profile as demonstrated in non-clinical studies with ARX788 [3].

Evaluating comparability of biologics due to manufacturing process changes from a quality perspective is based on the principles from ICH Q5E guidelines [7]. This assessment requires product-specific knowledge gathered through drug development to apply a totality-of-evidence approach to demonstrate comparability. Different levels of information are obtained from various studies including analytical studies for characterization of the molecule, animal studies for assessing toxicity, pharmacokinetics and pharmacodynamics, and clinical studies for assessing safety/tolerability, immunogenicity and efficacy. Analytical characterization of protein structure and function forms the foundation of comparability demonstration which can independently avoid conducting expensive and time-consuming animal and clinical studies [8–11].

Regulatory recommendation for ADC manufacturing involves rigorous control strategy and separate batch release of the cytotoxic payload and mAb intermediates, as well as the ADC drug substance (DS) and drug product (DP). Hence comparability studies need to assess the risk and impact of manufacturing changes on product quality for each of the intermediates, as well as the final conjugated bulk DS and DP. Depending on the structure of the payload, its solubility and clearance during DS manufacturing, the impact of any changes in manufacturing of the cytotoxic payload linker on product quality of DS and DP also needs to be assessed. Furthermore, confirmatory clinical use compatibility studies of the drug product with ancillary components including closed system transfer devices (CSTDs) used to prepare and deliver the ADC need to be performed [12].

The manufacturing process of ARX788 involves conjugation of ARX788 mAb to AS269 in a controlled reaction environment with continuous mixing, followed by diafiltration into formulation buffer and 0.2 micron filtration, to produce the ADC drug substance which is then stored frozen at -70°C. Drug Product manufacturing involves thawing the DS, mixing, sterile filtration and filling into glass vials in an Isolator. This is followed by washing of the DP vials, visual inspection, and storage at -20°C for the liquid formulation. For the new formulation and dosage form, the drug product vials are lyophilized and stored at 2-8°C. The manufacturing process of ARX788 DS and DP has undergone several changes during the life cycle from early to late-stage development which are summarized in Table 1. In this paper, we describe a comprehensive analytical comparability study between pre-change and post-change ARX788 materials to evaluate the impact of those changes, including manufacturing site change for DS and DP, and formulation and dosage form changes for DP. The mAb intermediate process did not undergo any significant changes, and thus did not warrant a separate comparability study. The AS269 intermediate underwent a minor change in manufacturing and was compared with pre-change lot using batch release and structural characterization methods. Further, ongoing long-term and accelerated GMP stability study results of ARX788 DS and DP were also monitored and the similarity of the stability trends were used to support the analytical comparability assessment (data not shown).

**Table 1:**
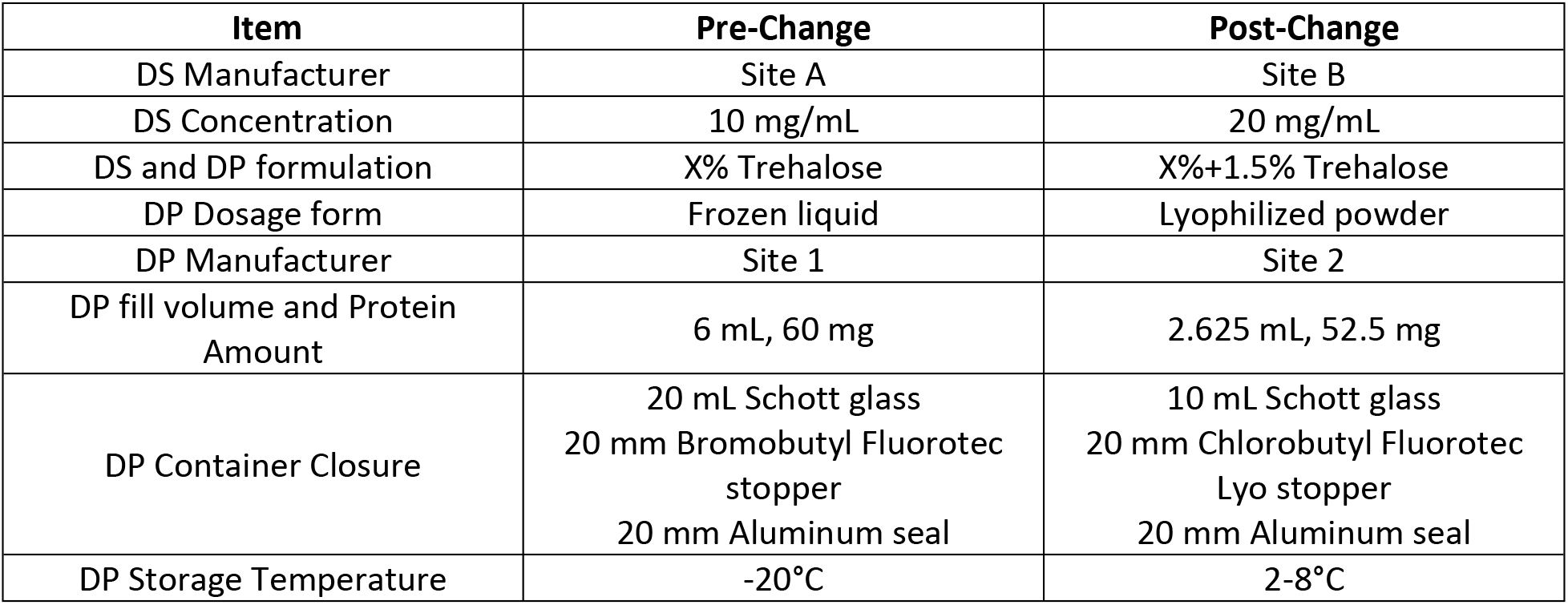
Summary of ARX788 manufacturing changes for DS and DP.

| Item | Pre-Change | Post-Change |
| --- | --- | --- |
| DS Manufacturer | Site A | Site B |
| DS Concentration | 10 mg/mL | 20 mg/mL |
| DS and DP formulation | X% Trehalose | X%+1.5% Trehalose |
| DP Dosage form | Frozen liquid | Lyophilized powder |
| DP Manufacturer | Site 1 | Site 2 |
| DP fill volume and Protein Amount | 6 mL, 60 mg | 2.625 mL, 52.5 mg |
| DP Container Closure | 20 mL Schott glass<br>20 mm Bromobutyl Fluorotec stopper<br>20 mm Aluminum seal | 10 mL Schott glass<br>20 mm Chlorobutyl Fluorotec Lyo stopper<br>20 mm Aluminum seal |
| DP Storage Temperature | -20°C | 2-8°C |

## Materials and Methods

### Ethics statement

No animal experiments were conducted in this study.

### Her2 Binding ELISA

An enzyme-linked immunosorbent assay (ELISA) was developed to measure ARX788 binding to target receptor HER2. Serially diluted ARX788 DS or DP samples were incubated in HER2-coated ELISA plates, and binding was detected with the addition of anti-human kappa-HRP antibody and incubation with 3,3’,5,5’-Tetramethylbenzidine (TMB) substrate. Sulfuric acid was added to stop the colorimetric reaction, and plates were read at absorbance wavelength 450 nm. The relative potency of pre-change or post-change ARX788 samples was calculated by comparison of its half-maximal effective concentration (EC50) value obtained from a 4-parameter sigmoidal fit of the dose response curve to the EC50 value of an ARX788 reference standard run on the same assay plate.

### HCC1954 Cytotoxicity Assay

The functional activity of ARX788 is measured with a cytotoxicity assay. HER2-positive HCC1954 breast cancer cells are incubated with serially diluted ARX788 DS or DP samples for 3 days, and cellular viability is quantified with addition of CCK-8 reagent. Viable cells cleave CCK-8 to a colored formazan dye, which is measured by absorbance at wavelength 450 nm. The relative potency of pre-change or post-change ARX788 samples was calculated by comparison of its half-maximal inhibitory concentration (IC_50_) value obtained from a 4-parameter sigmoidal fit of the dose response curve to the IC_50_ value of an ARX788 reference standard run on the same assay plate.

### SEC-HPLC

Size variants of ARX788 DS and DP batches were analyzed by Size Exclusion Chromatography (SEC) on a TSKgel G3000SWXL column (Tosoh Bioscience, catalog no.: 05841) using an Agilent 1100 or 1200 HPLC system with UV detection at 280 nm. Samples were injected at 50 ug load and separation was achieved by an isocratic elution using a mobile phase consisting of 200 mM potassium phosphate, 250 mM potassium chloride, pH 6.0 at a flow rate of 0.5 mL/min and column temperature set at 25°C. Percent area of HMW, monomer, and LWM species were calculated using Agilent ChemStation Software.

### CEX-HPLC

Cation Exchange chromatography (CEX) method was utilized to assess the charge variants of ARX788 DS and DP batches. Samples were loaded on a ProPac WCX-10, 4 x 250 mm column (Thermo Scientific) and eluted using a gradient of sodium chloride in a phosphate buffered mobile phase. Percent area of acidic, main, and basic species were calculated using Agilent ChemStation Software.

### HIC-HPLC

Hydrophobic interaction chromatography (HIC) method was utilized to assess the drug to antibody ratio (DAR) of ARX788 DS and DP batches. Samples were loaded on a Butyl-NPR HPLC column (Tosoh) and eluted using a gradient of ammonium sulphate in a phosphate buffered mobile phase. The drug to antibody ratio was determined by calculating the peak area of the mAb (DAR0), 1-drug (DAR1), and 2-drug (DAR2) forms using the following equation:

DAR = (% 1-drug + % 2-drug x 2)/(%mAb + % 1-drug + % 2-drug)

### Free Drug Related Impurities

ARX788 DS and DP batches were precipitated with acetonitrile in 2:3 ratio and vortexed for 1 minute. The samples were mixed on a thermomixer at room temperature and the mixtures were centrifuged for 30 minutes at 16,000 rcf. The supernatant was transferred to a new tube and mixed with hydroxylamine in a 1:1 ratio. The resulting solution was loaded to a XBridge Shield C-18 reverse phase column (Waters), followed by elution with a gradient of increasing acetonitrile content in mobile phase. The flow rate was set at 0.8 ml/min and detection wavelength was set at 214 nm. The amount of residual free drug of AS269 in the tested sample is determined by interpolation of observed peak area against a standard curve. The results are reported as a weight percent of ARX788 ADC.

### Intact mass determination

ARX788 DS samples were analyzed by LC-MS (Agilent 6200 Series TOF), following which molecular masses were obtained by deconvoluting the mass spectrum using Agilent MassHunter protein deconvolution software. For non-reduced mass analysis, no sample preparation was done prior to LC-MS analysis besides sample dilution in formulation buffer. For deglycosylation, the samples were treated with Peptide:N-Glycosidase F (PNGaseF) at 37°C for 2 hours prior to LC-MS analysis. For reduced mass analysis, the samples were treated with dithiothreitol (DTT) along with 4 M guanidine hydrochloride (GdnHCl) and heated at 70°C for 10 minutes before analysis by LC-MS.

All intact mass spectra were collected in positive ion mode with the mass range set to 700-3000 m/z with capillary and source temperatures at 300°C, sheath gas flow rate at 50 arbitrary units and auxiliary gas flow rate at 8 arbitrary units. The capillary voltage was set to 3500V with the fragmentor voltage set to 200V. The acquired spectra were deconvoluted using Agilent MassHunter protein deconvolution software.

### Peptide Mapping

Peptide maps of ARX788 DS and DP samples were obtained using an Reversed Phase Ultra Performance Liquid Chromatography (RP-UPLC) method with UV detection. Approximately 200 µg of samples were denatured in pH 8 Tris buffered 6M GdHCl, and reduced in 20 mM DTT by heating at 56°C for 30 min. The samples were allowed to cool to room temperature, and the reduced thiols were alkylated by incubation in 45 mM iodoacetamide in the dark for 30 min. The samples were then buffer exchanged using 0.5 mL Amicon Ultra 10 KDa MWCO spin-filters into trypsin digestion buffer containing 50mM TrisHCl pH 8, and 2M Urea. Sequencing grade Trypsin was then added, and the samples were incubated at 37°C for 4 hours. To stop the reaction, 15µL of 20% TFA was added to each sample.

Thereafter, 20µL volume of each sample were injected into a 4.6x150 mm Zorbax 300 SB-C8 (5µm) column and the peptides were separated using a gradient of TFA and acetonitrile at a flow rate of 1 mL/min and the UV spectra were acquired at 214 nm.

The disulfide bonds were mapped by comparing peptide maps of tryptic digests generated under non-reducing conditions.

For reduced peptide mapping, all samples were denatured, reduced, and alkylated, followed by digestion with Trypsin. The digests were analyzed by RP-UPLC with UV detection.

### icIEF

Imaged capillary iso-electric focusing (icIEF) was used to assess the pI and charge heterogeneity of ARX788 DS samples. Samples were prepared by adding and mixing appropriate amounts of carrier ampholytes, Urea, pI markers, arginine, and methylcellulose prior to analysis. The samples were loaded into the capillary and voltage applied, with pre-focusing condition at 1.5 kV for 2 minutes, and focusing condition at 3 kV for 8 minutes. Following the completion of focusing, a final image was taken at 280 nm for analysis. pI was calculated by comparing the position of the analyte to that of the pI standards. Percent main peak, acidic variants, and basic variants were also calculated based on area count of each species, relative to the total.

### N-Glycan Analysis

The N-linked glycoforms of ARX788 DS batches were enzymatically released using PNGase F, and fluorescently labeled using 2-amino benzamide (2-AB). Excess 2-AB dye was removed using acetone precipitation, and a Speedvac was used to remove residual acetone. The 2-AB labeled glycans were then reconstituted in water and the N-glycan profiles of the DS batches were obtained by hydrophilic interaction liquid chromatography (HILIC) UPLC analysis followed by fluorescent detection.

### Far UV Circular Dichroism

Far UV CD spectra (190 to 260 nm) of ARX788 DP lots were acquired using an Applied Photophysics Chirascan^TM^ Q100 Circular Dichroism Spectrometer containing a 0.01 cm cell at 20°C. All samples were diluted in formulation buffer to obtain a protein concentration of 0.5 mg/mL before analysis. After subtracting the baseline spectrum, the CD spectra of the samples were converted to mean residue molar ellipticity (molar ellipticity per residue) curves using the sample concentration, the mean residue weight, and the path-length of the cell.

### Near UV Circular Dichroism

ARX788 DP lots were diluted with formulation buffers to a concentration of 0.5 mg/mL before analysis. Near-UV (240 to 340 nm) CD measurements were carried out on an Applied Photophysics Chirascan^TM^ Q100 Circular Dichroism Spectrometer containing a 1 cm cell at 20°C. After subtracting the baseline spectrum, CD spectra of the samples were converted to mean residue molar ellipticity (molar ellipticity per residue) using the sample concentration, the mean residue weight, and the path-length of the cell.

### DSC

ARX788 DP lots were diluted to 1 mg/mL in formulation buffers and analyzed by MicroCal capillary VP Differential Scanning Calorimeter (DSC). The sample and reference cells were loaded with sample and formulation buffer, respectively. The instrument was programmed to scan from 10 to 110°C at a rate of 1°C/min, with a 10 second data averaging period and 15 min equilibration time. Buffer vs. buffer scans were recorded throughout the experiment sequence to obtain a baseline scan to subtract from the experimental data and to ensure the shapes of the baseline scans were reproducible over the course of the experiment. The raw data was processed using MicroCal PEAQ DSC software to calculate T_m_ values.

### SEC-MALS

Fifty micrograms of ARX788 DP were analyzed by SE-HPLC (Agilent 1100 HPLC system) on a TSKgel G3000SWXL column with inline Multi Angle Light Scattering (MALS) detector (Wyatt DAWN HELEOS) to determine the size distribution and MW of the monomer and HMW peaks. Separation on SEC column was achieved using an isocratic gradient with a mobile phase containing potassium phosphate and potassium chloride pH 6.0. Flow rate was set to 0.5 ml/min with column temperature set at 25°C. The SEC MALS data were analyzed using Astra v.7.3.2.

### SV-AUC

ARX788 DP lots and reference were diluted 20-fold using formulation buffer to a final concentration of 0.5 mg/ml before analysis of Sedimentation Velocity-Analytical Centrifugation (SV-AUC) using a Beckman Coulter XL-I analytical ultracentrifuge with double sector charcoal-filled Epon centerpiece. Protein sedimentation was accomplished through centrifugation at an angular velocity of 40,000 rpm. The concentration of each protein size variant was measured as a function of time and radial position using absorbance (A280) to monitor sedimentation. The concentration profiles were subsequently analyzed and plotted as a c(s) distribution.

### ADCC Effector Function

HER2-expressing SK-BR-3 cells were seeded at 5,000 cell/well in a 96-well assay plate, incubated overnight, and the growth medium was aspirated and replaced with 25 uL/well assay buffer (RPMI-1640+4% Low IgG Serum). Serially diluted ARX788 samples were added to the assay wells, followed by addition of 25 uL/well of Antibody Dependent Cellular Cytotoxicity (ADCC) Bioreporter Cells (Promega cat# G7010). The ADCC Bioreporter cells are engineered Jurkat cells stably expressing FcγRIIIa V158 (high affinity) receptors on the cell surface and an NFAT response element driving expression of firefly luciferase. After incubating assay plates for 6 hours at 37°C and 5% CO_2_, Bio-Glo Luciferase Assay Reagent was added, incubated for 30 minutes protected from light, and luminescence was measured on a SpectraMax plate reader.

The relative potency of ARX788 samples was calculated by comparison of its half-maximal effective concentration (EC_50_) values obtained from a 4-parameter sigmoidal fit of dose response curves against the EC_50_ value of the ARX788 reference standard run in the same assay plate.

### Fc Receptors Binding and FcRn Binding by Octet

The binding of ARX788 to Fc gamma receptors CD64 (FcγRI), CD32a (FcγRIIA) and CD16a (FcγRIIIA) was measured in a biolayer interferometry assay on an Octet RED96 system (Sartorius). Streptavidin-coated biosensors were loaded with purified biotinylated human CD64, CD32a, or CD16a in HBS-P+ buffer (Cytiva). After washing biosensors with HBS-P+ buffer to remove unbound protein, serially diluted ARX788 drug substance in HBS-P+ buffer was monitored for association and dissociation kinetics with loaded biosensors. Data were referenced using a parallel buffer blank subtraction. The processed binding curves were globally fitted using the Langmuir model describing a 1:1 binding stoichiometry. To avoid avidity effects in the CD32a and CD16a assays, the binding curves were globally fitted using 3.0 seconds for the dissociation step with the Langmuir model. Likewise, the FcRn binding assay was measured in a biolayer interferometry assay on an Octet RED96 system (Sartorius). Streptavidin-coated biosensors were loaded with purified biotinylated human FcRn in pH 6.0 assay buffer (100mM sodium phosphate, 150mM sodium chloride, 0.05% (v/v) Tween 20, pH 6.0). After washing biosensors with assay buffer to remove unbound protein, serially diluted ARX788 drug substance in pH 6.0 assay buffer was added and monitored for association and dissociation kinetics with FcRn loaded biosensors. Data were referenced using a parallel buffer blank subtraction. To avoid avidity effects, the processed binding curves were globally fitted using 3.0 seconds for the dissociation step with the Langmuir model describing a 1:1 binding stoichiometry.

## Results and discussions

### Side-by-side Batch Analysis

The batch release data of the pre-change and post-change DS and DP batches were compared side by side as shown in Table 2 and Table 3, respectively. For DS batches, all the purity assays including SEC-HPLC, CEX-HPLC, CE-SDS NR and R, and HIC-HPLC showed similar results. Likewise, the potency assays including HER2 Binding ELISA and HCC1954 Cytotoxicity Assay also showed similar results except for the first DS post-change batch ELISA at 125% and third post-change batch cytotoxicity at 74%, which is likely due to assay variability since the corresponding DP lot’s ELISA and cytotoxicity results are 94% and 103% (see Table 3), respectively. The protein concentration was increased from 10 mg/mL to 20 mg/mL in the post-change lots to reduce lyophilization cycle time during DP manufacturing. Trehalose concentration was also increased by 1.5% to help improve the stability of the drug product. These two changes accounted for the increase in osmolality from ∼157 to 210 mOsmol/kg. Likewise, the DP lot release analysis yielded similar results in most assays except for the expected differences in protein concentration and osmolality (Table 3). In summary, both pre- and post-change DS and DP batches passed the acceptance criteria for batch release and showed similar results.

**Table 2:** Bulk Drug Substance Batch Release Analysis of Pre-Change and Post-Change Batches.

| Tests |  | Pre-Change | Post-Change 1 | Post-Change 2 | Post-Change 3 |
| --- | --- | --- | --- | --- | --- |
| Peptide map |  | Conforms to RS | Conforms to RS | Conforms to RS | Conforms to RS |
| pH |  | 6.0 | 6.0 | 6.0 | 6.0 |
| Osmolality, mOsmol/kg |  | 157 | 210 | 209 | 209 |
| Protein concentration, mg/mL |  | 10.1 | 20.0 | 19.1 | 20.7 |
| HER2 Binding ELISA % relative to Reference |  | 98 | 125 | 117 | 107 |
| HCC1954 Cytotoxicity Assay, % relative to Reference |  | 106 | 96 | 90 | 74 |
| SEC-HPLC | Main % | 99.3 | 99.6 | 99.6 | 99.6 |
|  | HMW % | 0.6 | 0.4 | 0.4 | 0.4 |
|  | LMW % | 0.2 | ND | ND | ND |
| CEX-HPLC | Main % | 50.3 | 51.9 | 49.2 | 48.7 |
|  | Acidic % | 33.6 | 29.4 | 29.5 | 30.3 |
|  | Basic % | 16.1 | 18.6 | 21.3 | 21.0 |
| CE-SDS (NR) % purity |  | 95.1 | 96.9 | 97.7 | 98.1 |
| CE-SDS (R) % purity |  | 96.9 | 97.7 | 97.2 | 96.9 |
| HIC-HPLC | DAR | 1.9 | 1.9 | 1.9 | 1.9 |
|  | mAb | ND | 0.5 | ND | ND |
|  | Impurity | 5.4 | 3.8 | 4.2 | 4.0 |
| Free drug related impurities w/w |  | < 0.016% (LOQ) | 0.016% | 0.042% | 0.028% |

DAR = Drug antibody ratio; LOQ = limit of quantitation; NA = not tested; ND = not detected; RS= Reference Standard

**Table 3:**
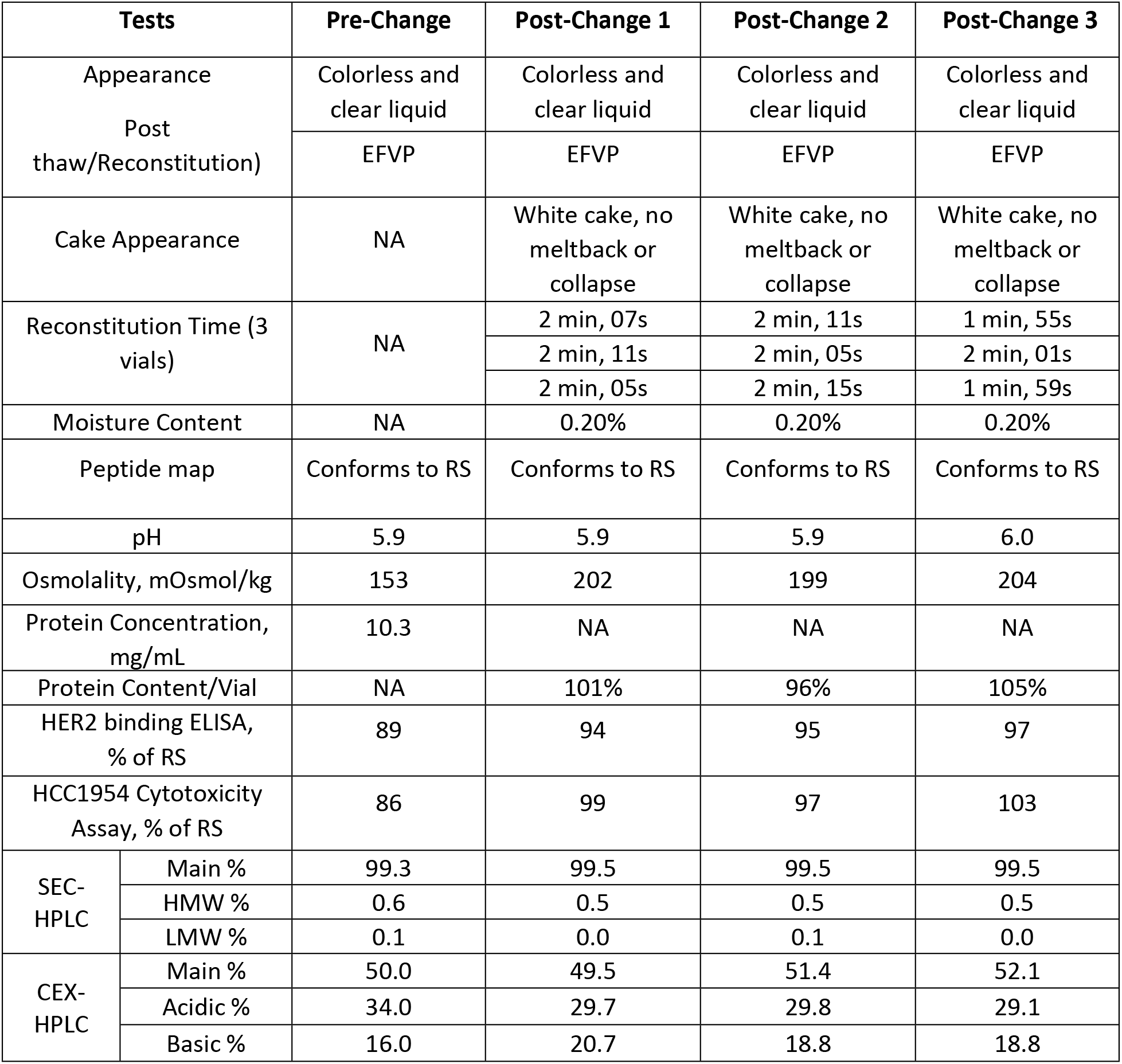
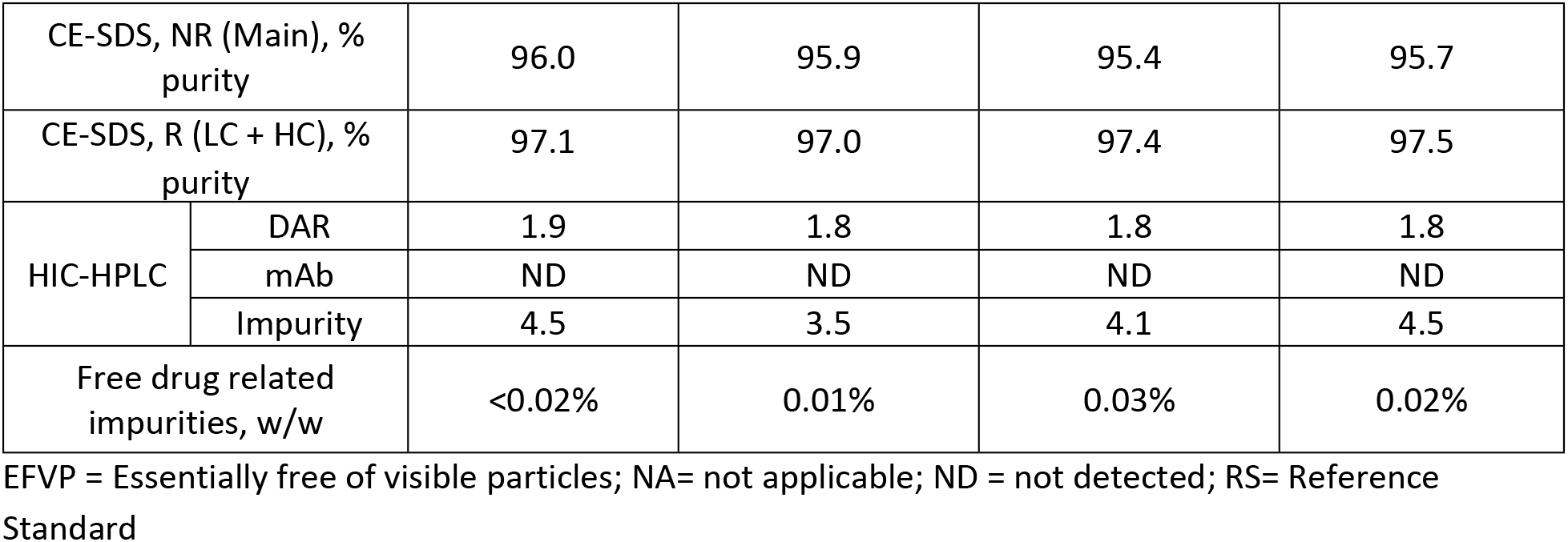
Drug Product Lot Release Analysis of Pre-Change and Post-Change Lots.

| Tests |  | Pre-Change | Post-Change 1 | Post-Change 2 | Post-Change 3 |
| --- | --- | --- | --- | --- | --- |
| Appearance<br>Post thaw/Reconstitution) |  | Colorless and clear liquid | Colorless and clear liquid | Colorless and clear liquid | Colorless and clear liquid |
|  |  | EFVP | EFVP | EFVP | EFVP |
| Cake Appearance |  | NA | White cake, no meltback or collapse | White cake, no meltback or collapse | White cake, no meltback or collapse |
| Reconstitution Time (3 vials) |  | NA | 2 min, 07s | 2 min, 11s | 1 min, 55s |
|  |  |  | 2 min, 11s | 2 min, 05s | 2 min, 01s |
|  |  |  | 2 min, 05s | 2 min, 15s | 1 min, 59s |
| Moisture Content |  | NA | 0.20% | 0.20% | 0.20% |
| Peptide map |  | Conforms to RS | Conforms to RS | Conforms to RS | Conforms to RS |
| pH |  | 5.9 | 5.9 | 5.9 | 6.0 |
| Osmolality, mOsmol/kg |  | 153 | 202 | 199 | 204 |
| Protein Concentration, mg/mL |  | 10.3 | NA | NA | NA |
| Protein Content/Vial |  | NA | 101% | 96% | 105% |
| HER2 binding ELISA, % of RS |  | 89 | 94 | 95 | 97 |
| HCC1954 Cytotoxicity Assay, % of RS |  | 86 | 99 | 97 | 103 |
| SEC-HPLC | Main % | 99.3 | 99.5 | 99.5 | 99.5 |
|  | HMW % | 0.6 | 0.5 | 0.5 | 0.5 |
|  | LMW % | 0.1 | 0.0 | 0.1 | 0.0 |
| CEX-HPLC | Main % | 50.0 | 49.5 | 51.4 | 52.1 |
|  | Acidic % | 34.0 | 29.7 | 29.8 | 29.1 |
|  | Basic % | 16.0 | 20.7 | 18.8 | 18.8 |
| CE-SDS, NR (Main), % purity |  | 96.0 | 95.9 | 95.4 | 95.7 |
| CE-SDS, R (LC + HC), % purity |  | 97.1 | 97.0 | 97.4 | 97.5 |
| HIC-HPLC | DAR | 1.9 | 1.8 | 1.8 | 1.8 |
|  | mAb | ND | ND | ND | ND |
|  | Impurity | 4.5 | 3.5 | 4.1 | 4.5 |
| Free drug related impurities, w/w |  | <0.02% | 0.01% | 0.03% | 0.02% |

EFVP = Essentially free of visible particles; NA= not applicable; ND = not detected; RS= Reference Standard

### Side-by-Side Physicochemical Analysis

A panel of physicochemical tests were conducted for both pre- and post-change BDS batches including intact mass analysis under both Non-reducing and reducing conditions, peptide mapping, icIEF, and N-glycan analysis. Analytical methods and target acceptance criteria for assessing product comparability between pre-change and post-change DS and DP batches manufactured for clinical development were pre-determined before the execution of the study.

### Intact Mass Analysis, Non-reduced

ARX788 DS was analyzed by LC-MS (Agilent 6200 Series TOF), following which molecular masses were obtained by deconvoluting the mass spectrum using Agilent Protein Deconvolution software. The deconvoluted spectra of DS obtained from pre-change and post-change1 materials as well as the Reference Standard (RS) are shown in Fig 1A. The measured molecular masses showing different glycan forms are listed in Table 4. The molecular masses of all batches of DS are consistent with the theoretical values and the similarity of results indicate that all batches are comparable.

**Table 4:** Molecular Mass of ARX788 Drug Substance.

| Sample | Modifier | Measured Mass (Da) | Mass Difference relative to Pre-Change Sample (Da) |
| --- | --- | --- | --- |
| Pre-Change | G0F/G0F | 150049.89 | NA |
|  | G0F/G1F | 150206.23 | NA |
|  | G1F/G1F or G0F/G2F | 150367.56 | NA |
| Post-Change 1 | G0F/G0F | 150060.08 | 10.19 |
|  | G0F/G1F | 150220.66 | 14.43 |
|  | G1F/G1F or G0F/G2F | 150375.91 | 8.35 |
| Post-Change 2 | G0F/G0F | 150058.12 | 8.23 |
|  | G0F/G1F | 150219.68 | 13.45 |
|  | G1F/G1F or G0F/G2F | 150375.53 | 7.97 |
| Post-Change 3 | G0F/G0F | 150062.02 | 12.13 |
|  | G0F/G1F | 150222.69 | 16.46 |
|  | G1F/G1F or G0F/G2F | 150374.61 | 7.05 |
| Reference Standard | G0F/G0F | 150047.56 | -2.33 |
|  | G0F/G1F | 150215.37 | 9.14 |
|  | G1F/G1F or G0F/G2F | 150368.29 | 0.73 |

The deconvoluted spectra of reduced DS batches obtained from pre-change and post-change are shown in Fig 1B for heavy chain. The measured molecular masses of light chain and heavy chain observed in both batches are listed in Table 5. The results are consistent with the theoretical values and the similarity of results indicate the batches are comparable.

**Table 5:** Molecular Mass of Reduced ARX788 Drug Substance.

| Sample | Modifier | Measured Mass (Da) | Mass Difference relative to Pre-Change Sample (Da) |
| --- | --- | --- | --- |
| Pre-Change | LC | 23444.20 | NA |
|  | HC:G0F | 51595.67 | NA |
|  | HC:G1F | 51757.83 | NA |
|  | HC:G2F | 51918.57 | NA |
| Post-Change 1 | LC | 23444.28 | 0.08 |
|  | HC:G0F | 51595.80 | 0.13 |
|  | HC:G1F | 51758.00 | 0.17 |
|  | HC:G2F | 51919.05 | 0.48 |
| Post-Change 2 | LC | 23444.23 | 0.03 |
|  | HC:G0F | 51595.73 | 0.06 |
|  | HC:G1F | 51757.79 | -0.04 |
|  | HC:G2F | 51918.51 | -0.06 |
| Post-Change 3 | LC | 23444.28 | 0.08 |
|  | HC:G0F | 51595.95 | 0.28 |
|  | HC:G1F | 51758.09 | 0.26 |
|  | HC:G2F | 51918.43 | -0.14 |
| Reference Standard | LC | 23444.05 | 0.08 |
|  | HC:G0F | 51595.49 | 0.13 |
|  | HC:G1F | 51757.60 | 0.17 |
|  | HC:G2F | 51918.00 | 0.48 |

### Peptide Mapping

ARX788 DS samples were analyzed by peptide mapping to confirm the comparability of their primary sequences. All samples were denatured, reduced, and alkylated, followed by digestion with Trypsin. The digests were analyzed by RPHPLC with UV 214 detection. The peptides were well resolved and provided comparable UV profiles of pre-change and post-change1 as shown in Fig 1C. The para-acetyl phenylalanine (pAF) and AS269 containing peptide highlighted in the chromatograms was also observed in all batches with comparable peak intensities. These results indicate that all batches of ARX788 DS are chemically comparable.

**Figure 1:**
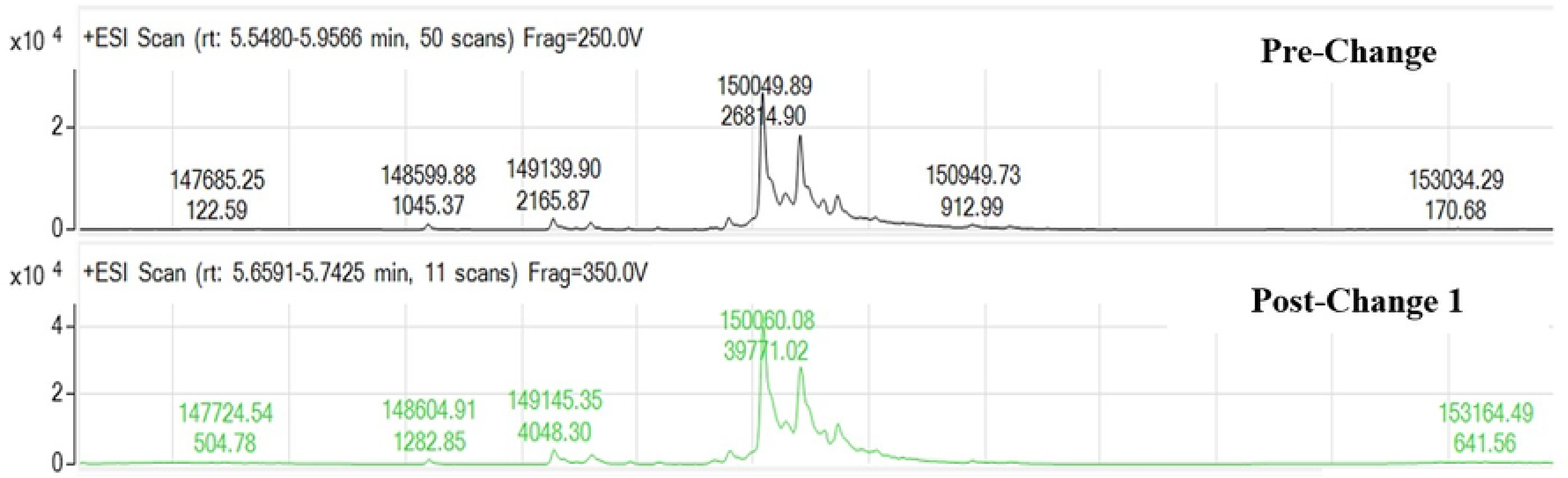
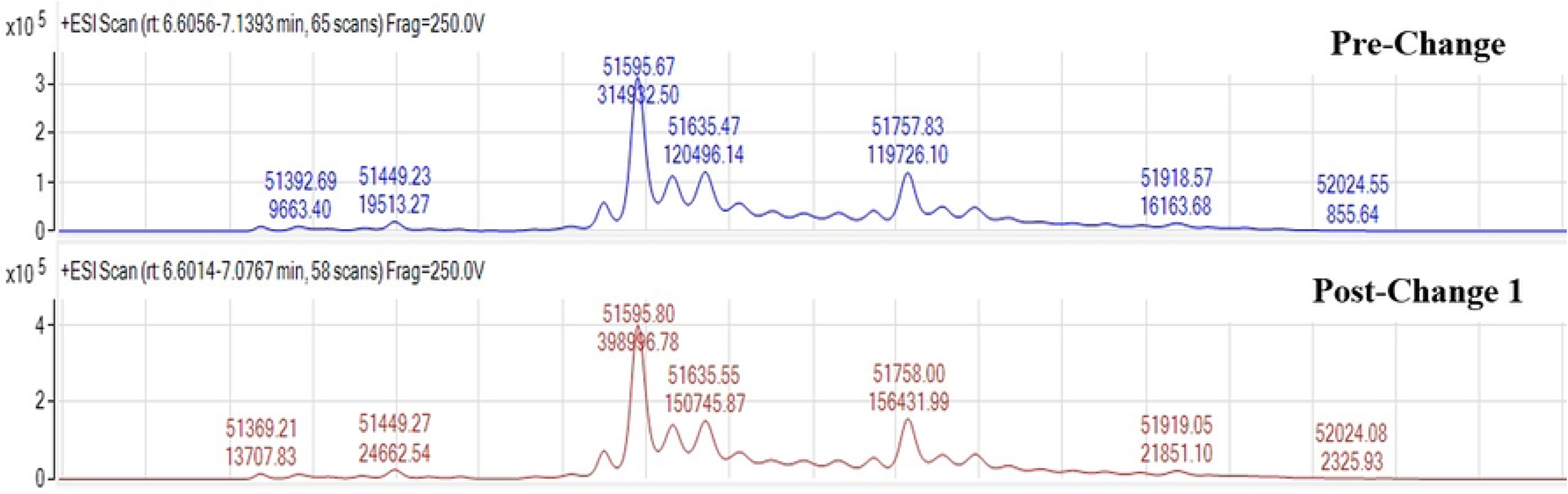
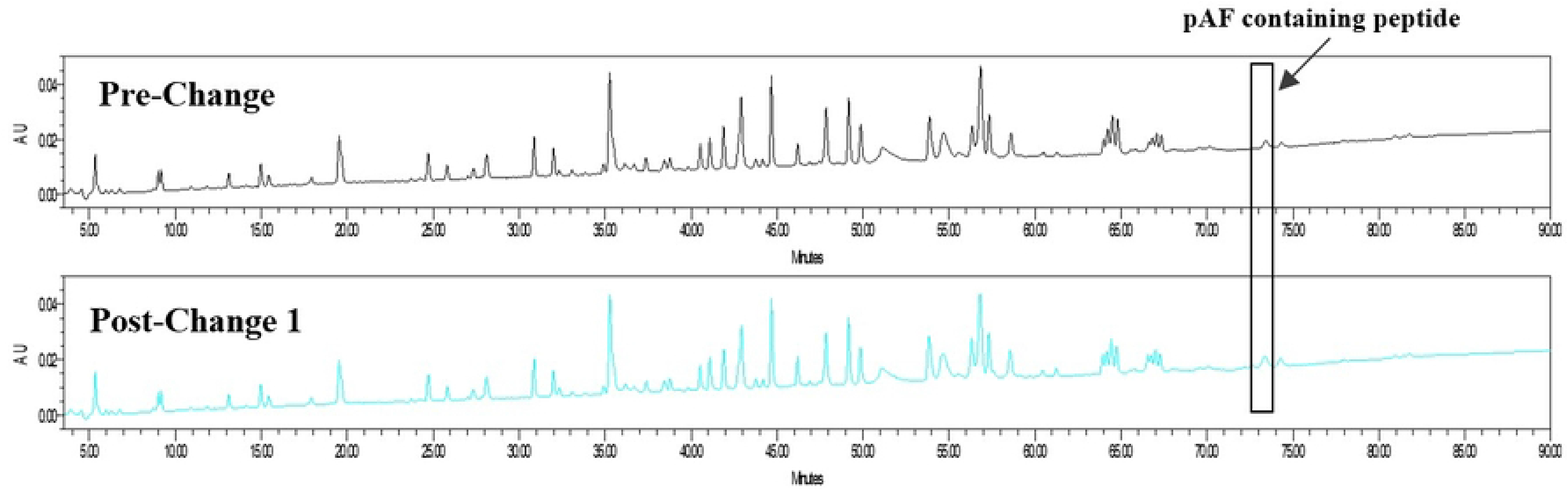
A) Non-reduced Deconvoluted Mass Spectra of ARX788 Drug Substance Batches, B) Reduced Deconvoluted Mass Spectra of Heavy Chains in ARX788 Drug Substance, and C) 214 nm UV Profiles of ARX788 Drug Substance Trypsin Digest Peptide Map.

### Imaged capillary isoelectric focusing (icIEF)

icIEF electrophoresis method separates proteins based on differences in isoelectic point (pI) using a pH gradient formed from ampholytes in an electric field. The pI value of the protein is determined by comparing against two pI standard markers. ARX788 DS batches from pre-change and post-change were analyzed using ProteinSimple Imaged icIEF system with a fluorocarbon-coated electrophoresis capillary for the determination of isoelectric point (pI) and detection of charge variants. As shown in Fig 2A, the UV profiles of all batches of ARX788 DS were highly similar and displayed a main peak pI value of 8.4. The relative percentage of charge variants were also similar among the batches (Table 6) with 0.7% difference in main peak purity between Pre-change and post-Change2 batches. Overall, these results indicate a high level of comparability among the batches of ARX788 DS.

**Table 6:** Isoelectric Point and Relative Abundance of Charge Variants in ARX788 Dug Substance Determined by icIEF

| Sample | pI | % Acidic | %Main | % Basic |
| --- | --- | --- | --- | --- |
| Reference Standard | 8.4 | 42.5 | 49.1 | 8.4 |
| Pre-Change | 8.4 | 43.4 | 48.7 | 7.9 |
| Post-Change 1 | 8.4 | 39.4 | 49.3 | 11.3 |
| Post-Change 2 | 8.4 | 39.7 | 49.4 | 11.0 |
| Post-Change 3 | 8.4 | 41.2 | 49.2 | 9.6 |

### N-Glycan Analysis

The N-linked glycoforms of ARX788 DS batches were enzymatically released using PNGase F and fluorescently labeled using 2-amino benzamide (2-AB). The N-glycan profiles of the DS batches were obtained by using hydrophilic interaction liquid chromatography (HILIC) UPLC analysis and fluorescent detection. The N-glycan profiles for all four batches of DS and reference are shown in Fig 2B and the results are summarized in Table 7**Error! Reference source not found.**. The results show that the most abundant N-glycan forms are G0, G0F, Man-5, G1, G1F, and G2F and the glycan profiles are almost identical indicating a high degree of comparability among the DS batches from pre-change and post-change.

**Figure 2:**
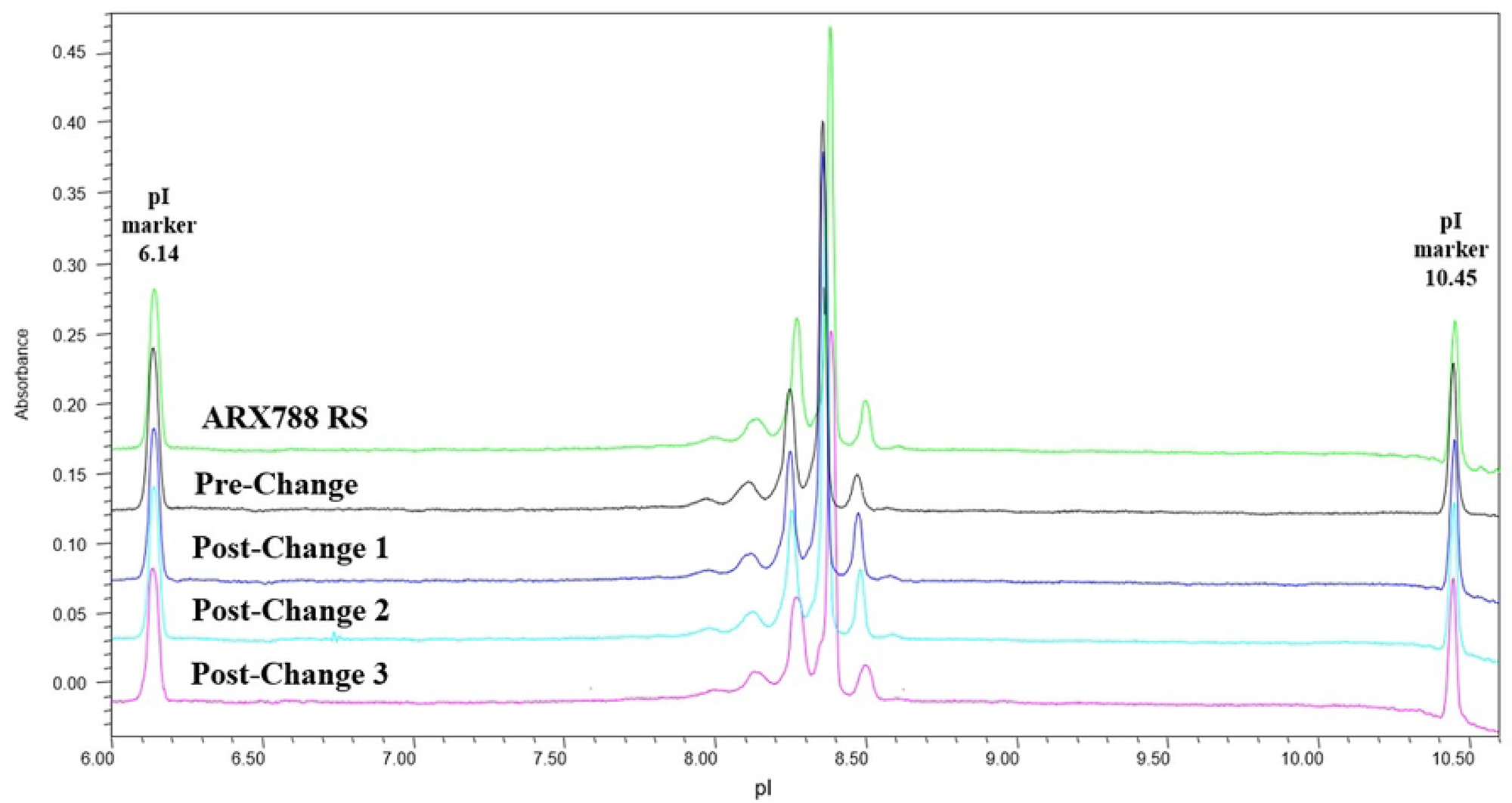
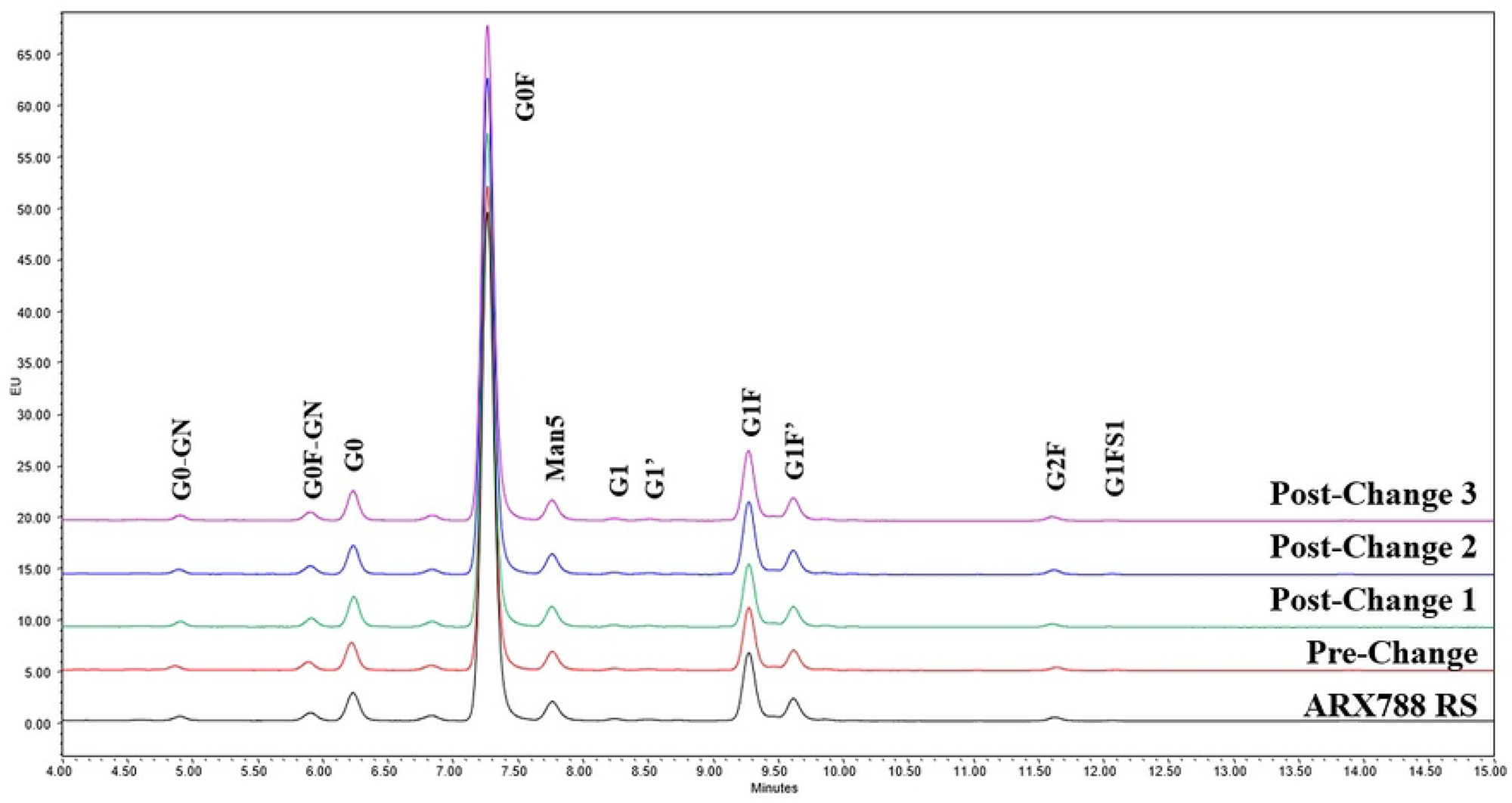
A) Imaged Capillary Isoelectric Focusing Electropherograms and B) HILIC-FL Profile of 2-AB Labeled N-Glycans from ARX788 DS batches

**Table 7:**
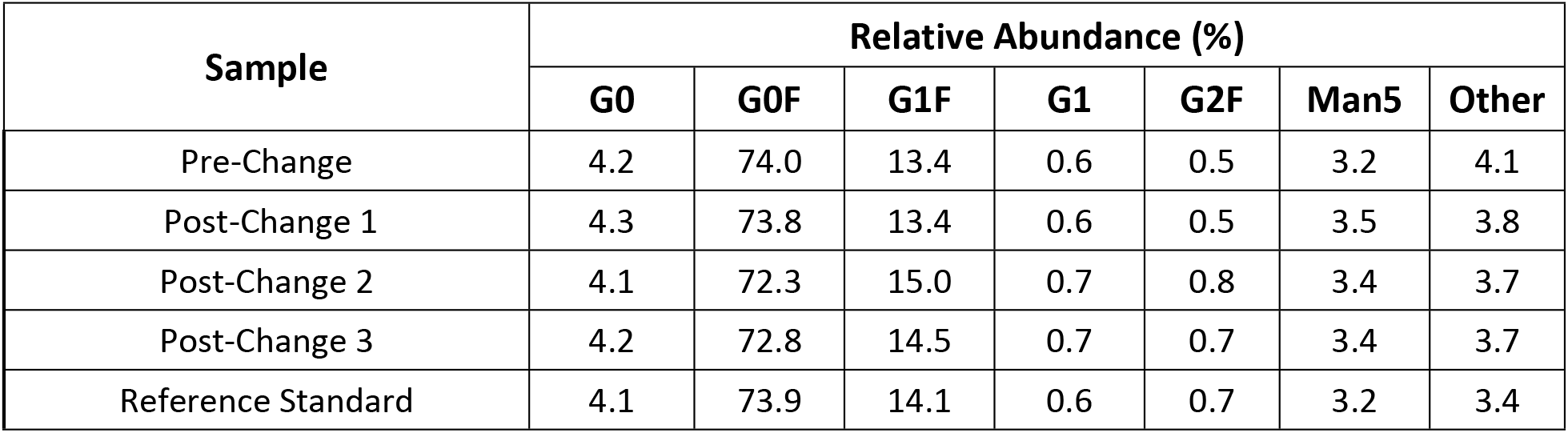
Relative Abundance of N-Glycan Forms in ARX788 Drug Substance.

| Sample | Relative Abundance (%) |  |  |  |  |  |  |
| --- | --- | --- | --- | --- | --- | --- | --- |
|  | G0 | G0F | G1F | G1 | G2F | Man5 | Other |
| Pre-Change | 4.2 | 74.0 | 13.4 | 0.6 | 0.5 | 3.2 | 4.1 |
| Post-Change 1 | 4.3 | 73.8 | 13.4 | 0.6 | 0.5 | 3.5 | 3.8 |
| Post-Change 2 | 4.1 | 72.3 | 15.0 | 0.7 | 0.8 | 3.4 | 3.7 |
| Post-Change 3 | 4.2 | 72.8 | 14.5 | 0.7 | 0.7 | 3.4 | 3.7 |
| Reference Standard | 4.1 | 73.9 | 14.1 | 0.6 | 0.7 | 3.2 | 3.4 |

### Side-by-Side Biophysical Characterization

Various biophysical tests were conducted for both pre- and post-change DP lots. Far UV CD was employed for secondary structure comparison while near UV CD and DSC were carried out for tertiary structure comparison. The size distributions of ARX788 DP lots were compared using two different biophysical techniques, namely SEC-MALS, and SV-AUC.

### Secondary Structural Comparison

Far UV CD was employed to assess the secondary structure element of these samples. The overlaid mean residue molar ellipticity (MRME) results are presented in Fig 3A and the spectra are identical within experimental error, which shows that the relative composition of secondary structural elements are highly comparable. These results demonstrate a high degree of spectral similarity indicating lot-to-lot consistency of secondary structure.

### Tertiary Structural Comparison

The near UV CD determines the tertiary structure due to asymmetric environments of tryptophan, tyrosine, phenylalanine, and disulfide. The spectra are shown in Fig 3B, which shows a high degree of similarity between pre-change and post-change drug product lots demonstrating their comparability at the level of tertiary structure. Likewise, Capillary differential scanning calorimetry (DSC) was employed to detect the thermal transition temperature (Tm) of proteins for thermal stability determination. Proteins with different thermal stabilities show different Tm values that reflect their higher order folded structures. The DSC thermograms of the drug product lots along with reference are shown in Fig 3C which shows that the peak shape and transition temperatures are nearly identical. The results demonstrate that pre-change and post-change Lyo ARX788 DP materials have the same Tm values and are therefore comparable.

### Size Comparison by SEC-MALS

ARX788 DP lots were analyzed by SE-HPLC with inline MALS detector to determine the size distribution and MW of the monomer and HMW peaks. The results are shown in Fig 3D. All four batches of ARX788 drug product show identical 153-155kD MW for the monomer peak. The MW of HMW species in these samples also correlate roughly with that of a dimer of the main peak considering higher variability of MW determination because of low abundance (signal) of the HMW peak.

### Size Comparison by SV-AUC

Sedimentation velocity (SV) measurement by analytical ultracentrifugation (AUC) is a classical method for protein size analysis and state of association or aggregation. Aggregates can be detected based on difference in sedimentation coefficients which is a function of shape and mass of the oligomers. ARX788 DP lots were compared using SV-AUC and the resulting plot of c(s) distribution is presented in Fig 3E. The sedimentation coefficients of the main peak are slightly different between pre-change and the 3 lots of post-change DP. This difference is due to the fact that the formulation buffer of the latter samples contains slightly higher Trehalose concentration, which leads to an increase in viscosity and density, thus lowering the sedimentation rate of post-change samples. The data also suggest that pre-change may contains slightly higher HMW species although it is difficult to quantitatively assess the difference as the values were below detection threshold of 3.7% (note that SE-HPLC as well as SEC-MALS show very similar HMW levels). Based on these results, it is concluded that theses samples have comparable size distributions by SV-AUC, showing >96% main peak and ≤3.1% HMW species for all DP lots.

**Figure 3:**
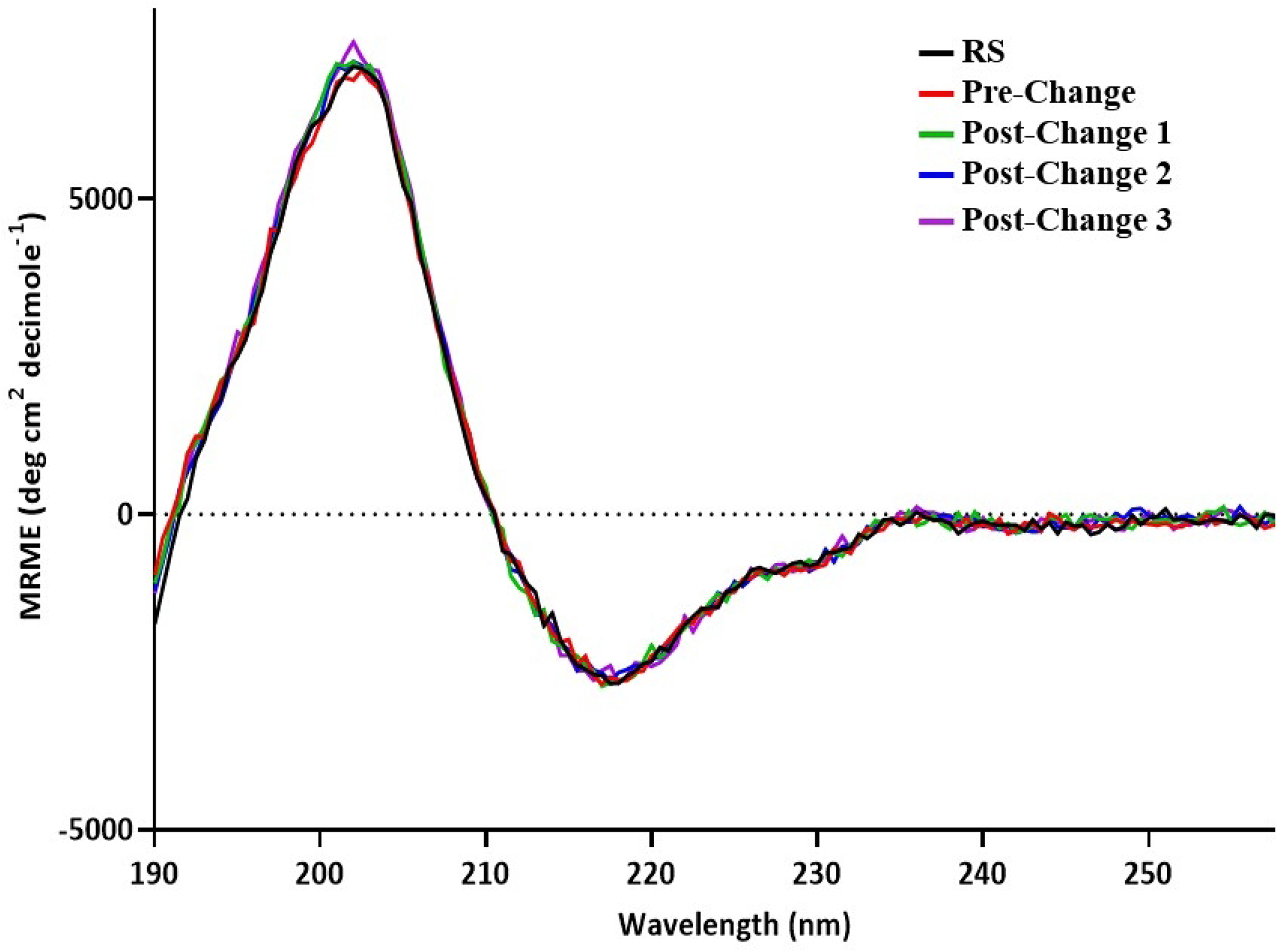
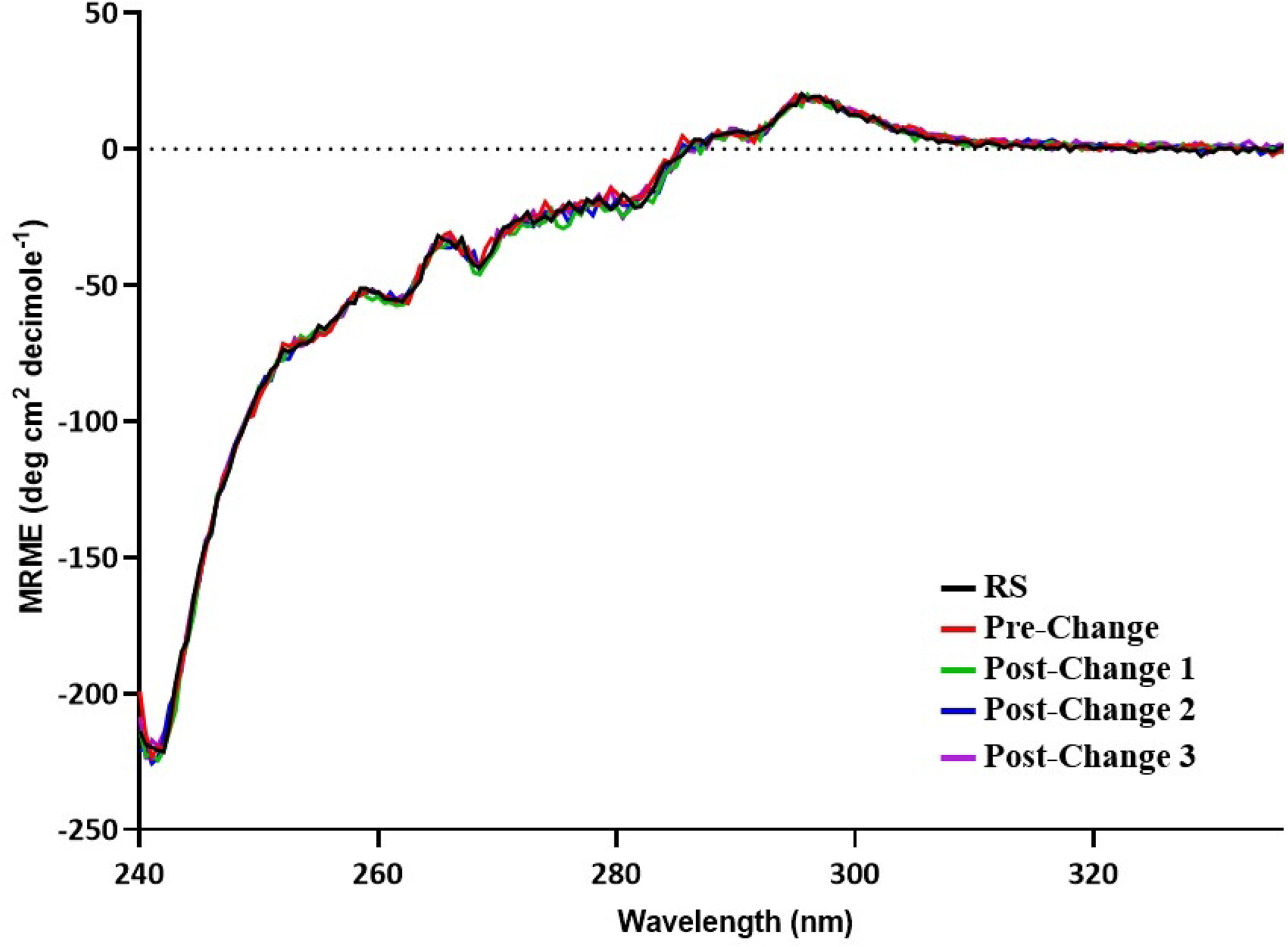
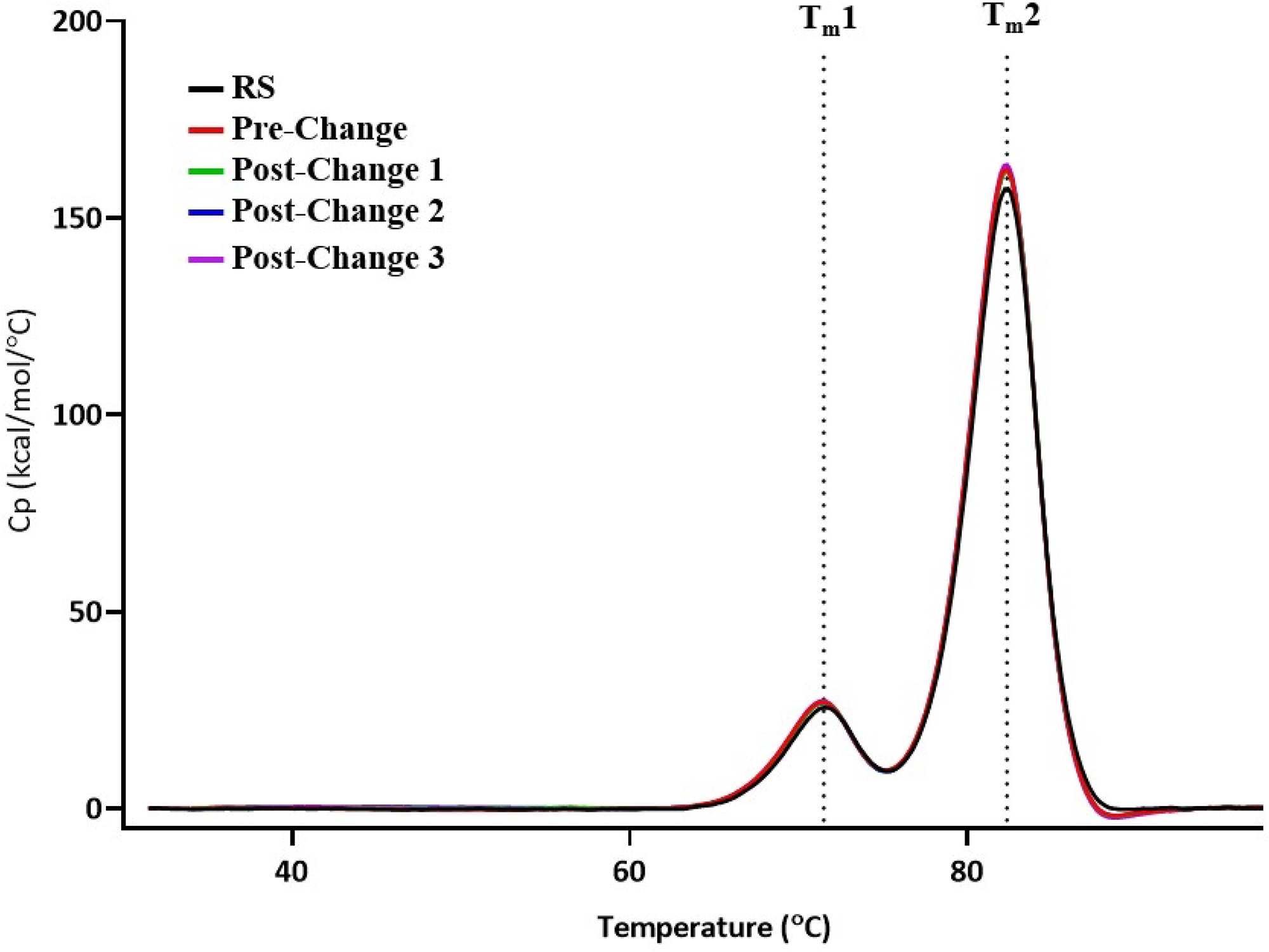
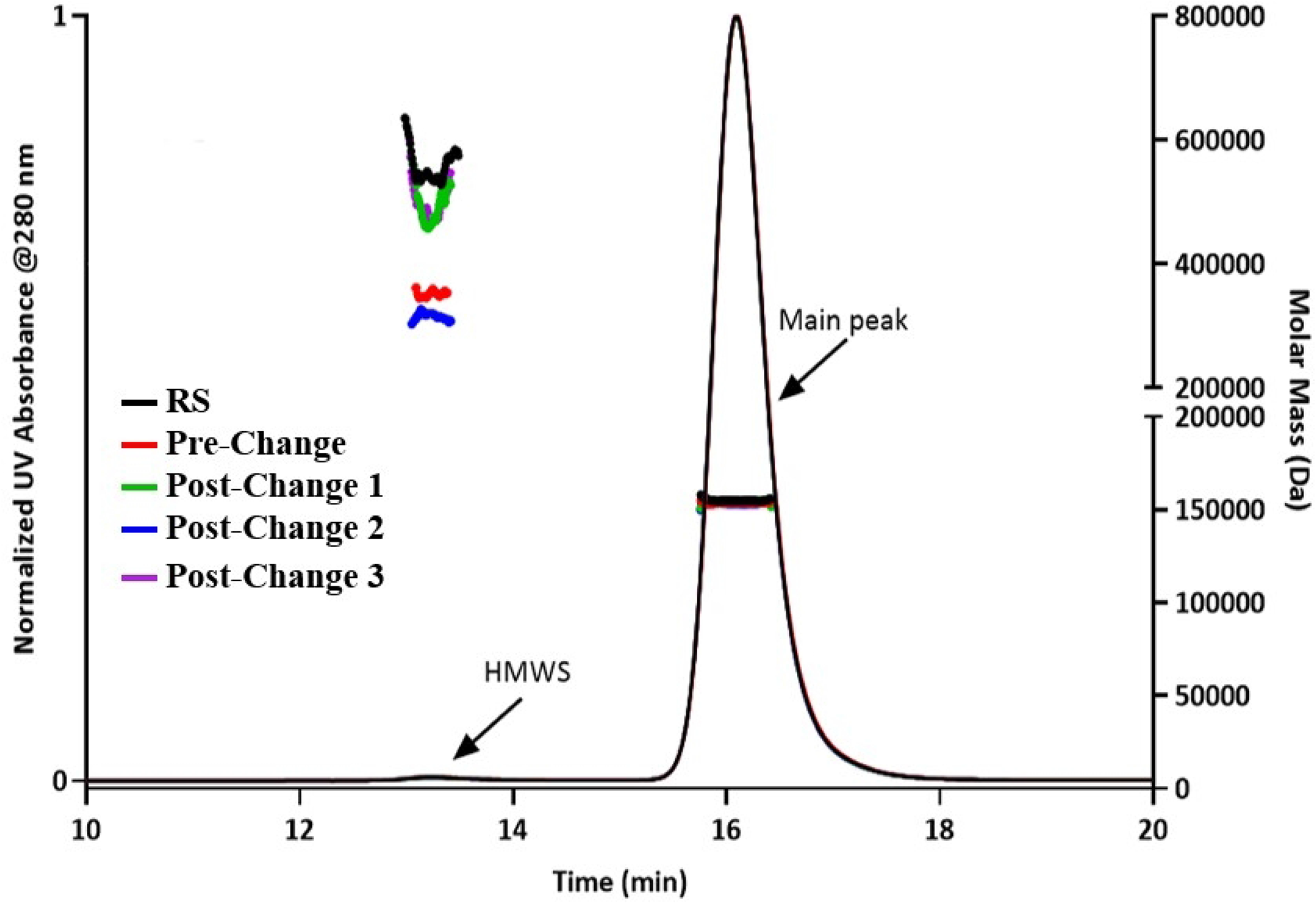
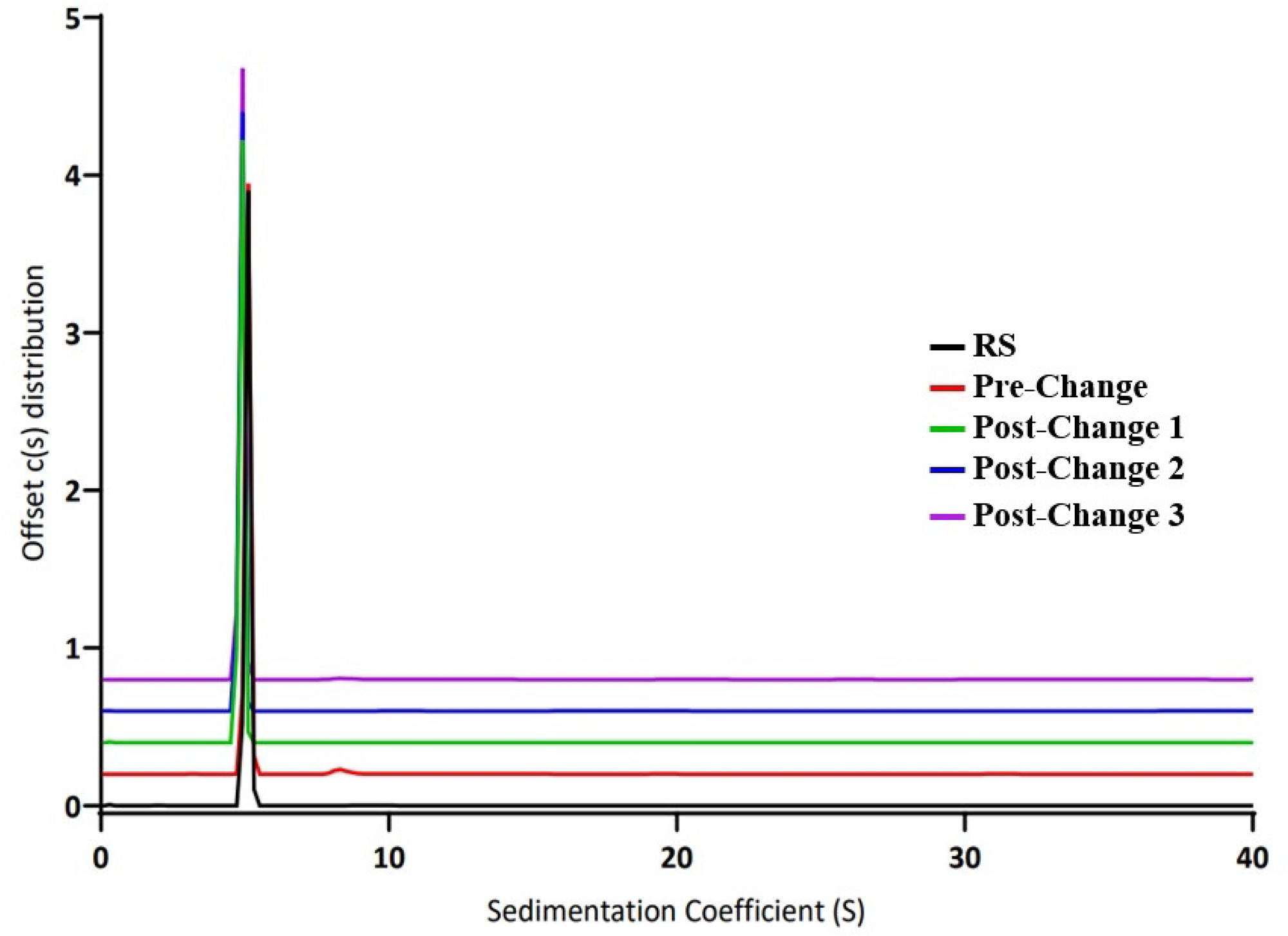
Biophysical Characterization of ARX788 DP lots: A) Far UV CD spectra, B) Near UV CD spectra, D) DSC thermograms, D) SEC-MALS chromatograms, and E) SV-AUC c(s) distribution plot.

### Side-by-Side Biological Characterization

#### ADCC Functional Activity Comparison

Although not the primary mechanism-of-action of ARX788, ADCC activity of pre-change and post-change batches were measured in an ADCC reporter assay Representative dose response curves for ARX788 reference standard, DS Pre-change batch, and DS Post-change batches 1 and 2, are shown in Fig 4A and the average Relative Potency and fold difference values are summarized in Table 8. The 1.17 to 1.42-fold-difference in Relative Potency of ARX788 DS post-change versus pre-change batches fell within the variability of the assay and met the defined difference in relative potency criterion. Thus, the observed ADCC activity is comparable between ARX788 DS pre-change and post-change batches.

**Table 8:**
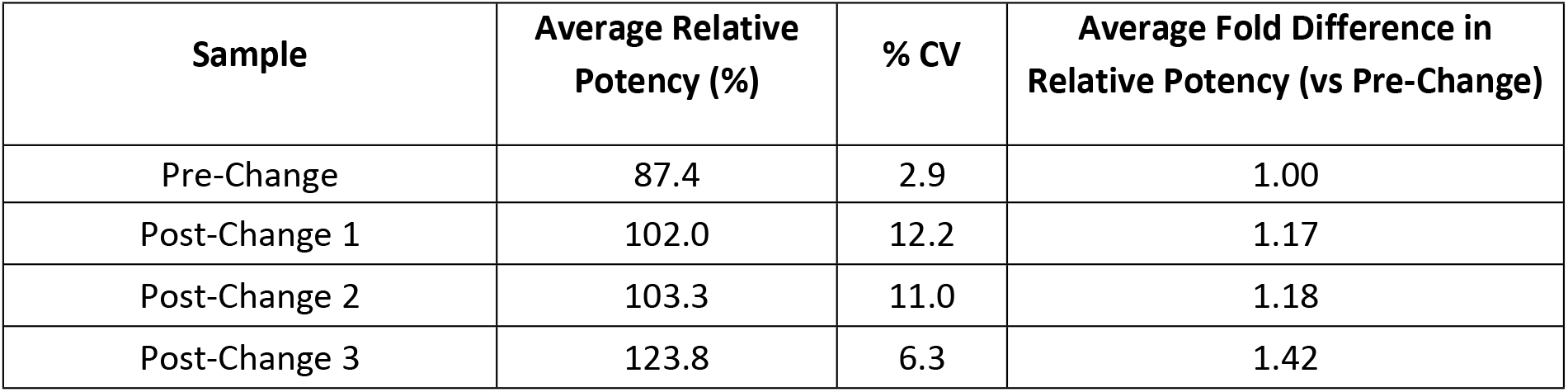
Average Fold Difference in Relative ADCC Potency of ARX788 DS Batches

#### Fc Receptor Affinity Comparison

An orthogonal method for analyzing effector function is by examining the binding affinity of ARX788 from pre- and post-change batches towards Fc gamma receptors including CD64 (FcγR I), CD32a (FcγR IIA), CD16a (FcγR IIIA). The binding kinetics sensorgrams of pre-change and post-change1 DS to the three Fc gamma receptors are shown in Fig4B and the dissociation constants (KD) of all samples are summarized in Table 9. Pre-change and post-change DS samples showed highly similar binding kinetics for the three tested Fc gamma receptors as seen in the sensorgrams, which suggested that these samples have similar affinities towards the receptors. Indeed, the KD values are highly similar between RS to pre-change as well as RS to post-change batches (KD ratios are very close to 1.00). Thus, the Fc gamma binding characteristics are comparable among the four DS batches.

**Table 9.** : Fc Gamma and FcRn Receptors Binding Affinities for ARX788 DS Batches

| Sample | CD64 K<sub>D</sub> Ratio | CD32a K<sub>D</sub> ratio (Sample/RS) | CD16a K<sub>D</sub> Ratio (Sample/RS) | FcRn K<sub>D</sub> Ratio (Sample/RS) |
| --- | --- | --- | --- | --- |
| Reference | NA | NA | NA | 1.00 |
| Pre-Change | 1.00 | 1.28 | 0.99 | 0.98 |
| Post-Change 1 | 1.05 | 1.25 | 1.13 | 1.60 |
| Post-Change 2 | 0.98 | 1.22 | 1.03 | 1.33 |
| Post-Change 3 | 0.89 | 1.05 | 1.01 | 1.19 |

Analysis of FcRn binding by ARX788 should also provide insight into effector function comparability. The binding kinetics sensorgrams of pre-change and post-change1 DS to FcRn receptor are shown in Figure 4C and the dissociation constants (KD) of all samples are summarized in Table 9. All pre-change and post-change DS samples showed highly similar binding kinetics for FcRn receptor as seen in the sensorgrams, which suggested that these samples have similar affinities towards the receptors. Indeed, the KD values are highly similar between RS to pre-change as well as RS to post-change batches (KD ratios very close to 1). Thus, the FcRn binding affinities are comparable among the four DS batches.

**Figure 4:**
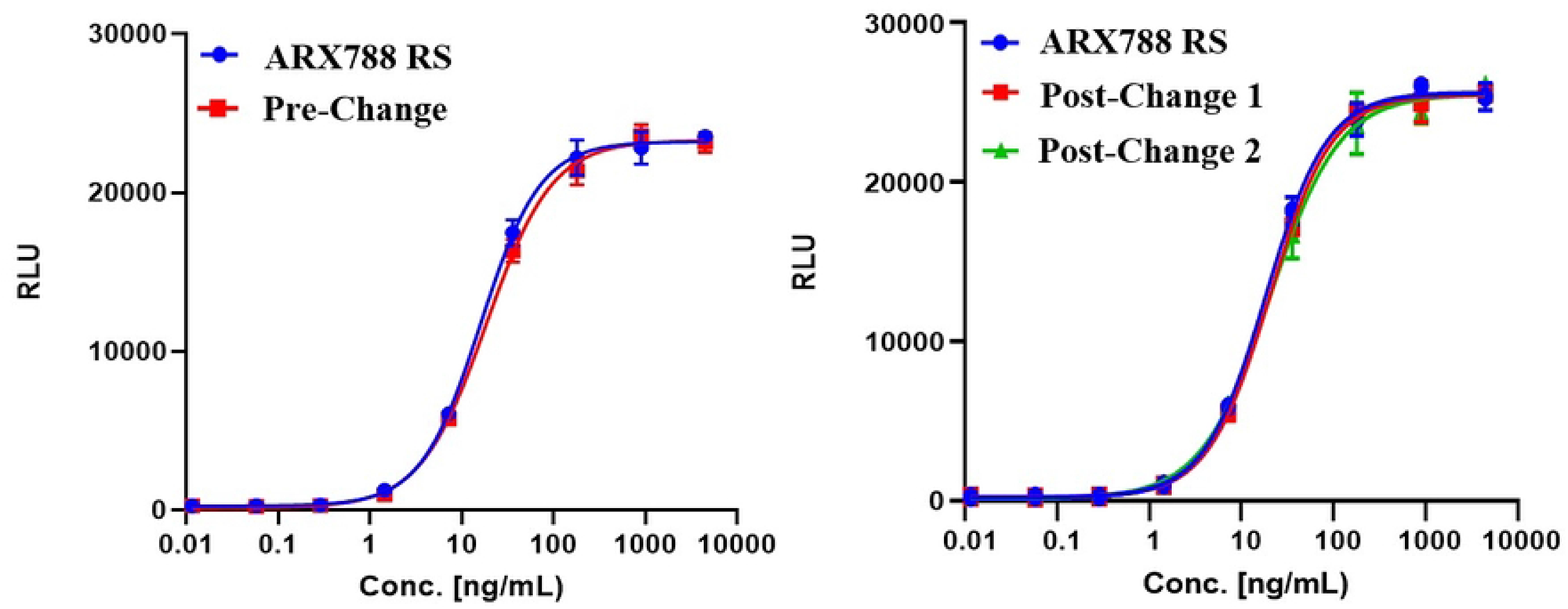
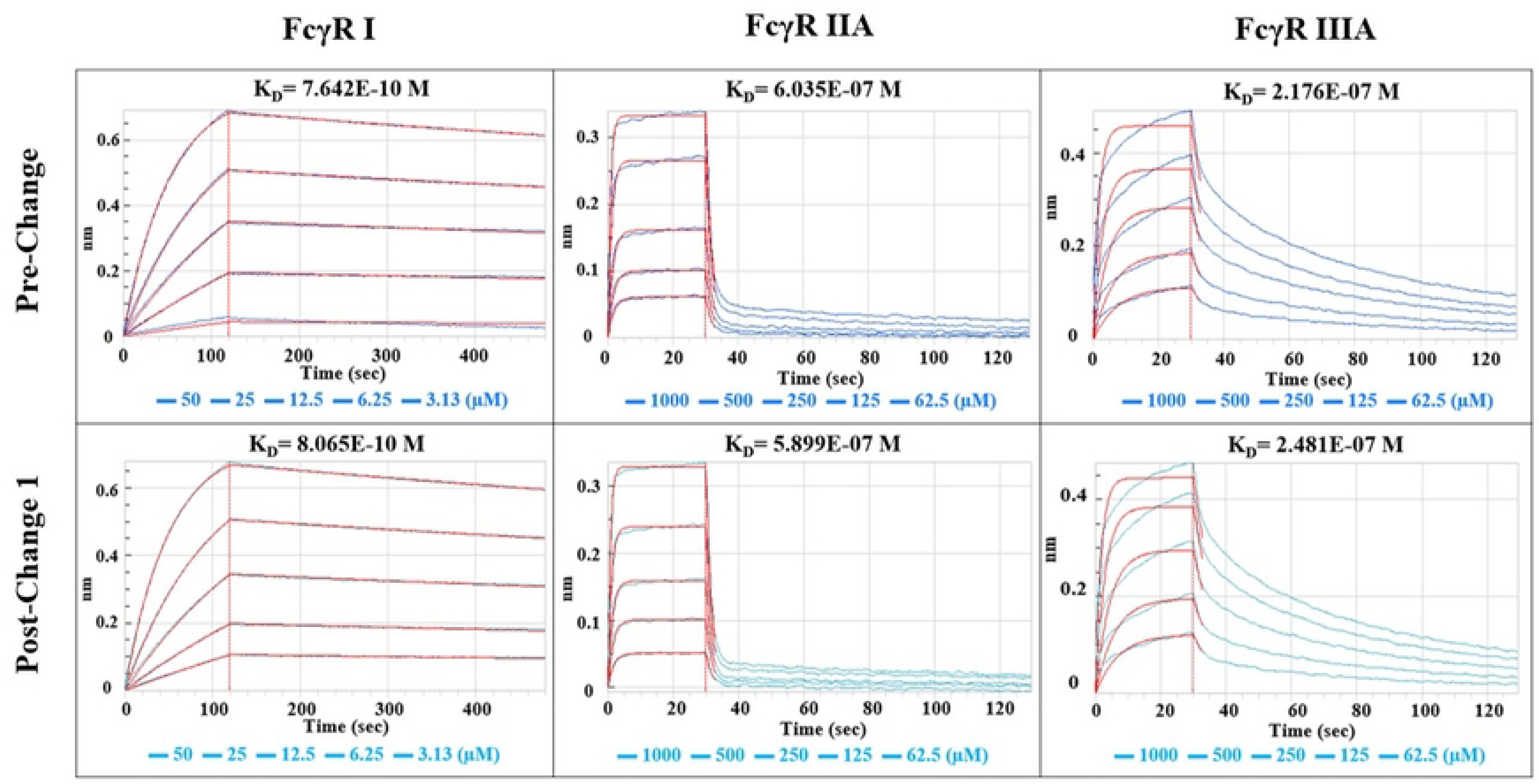
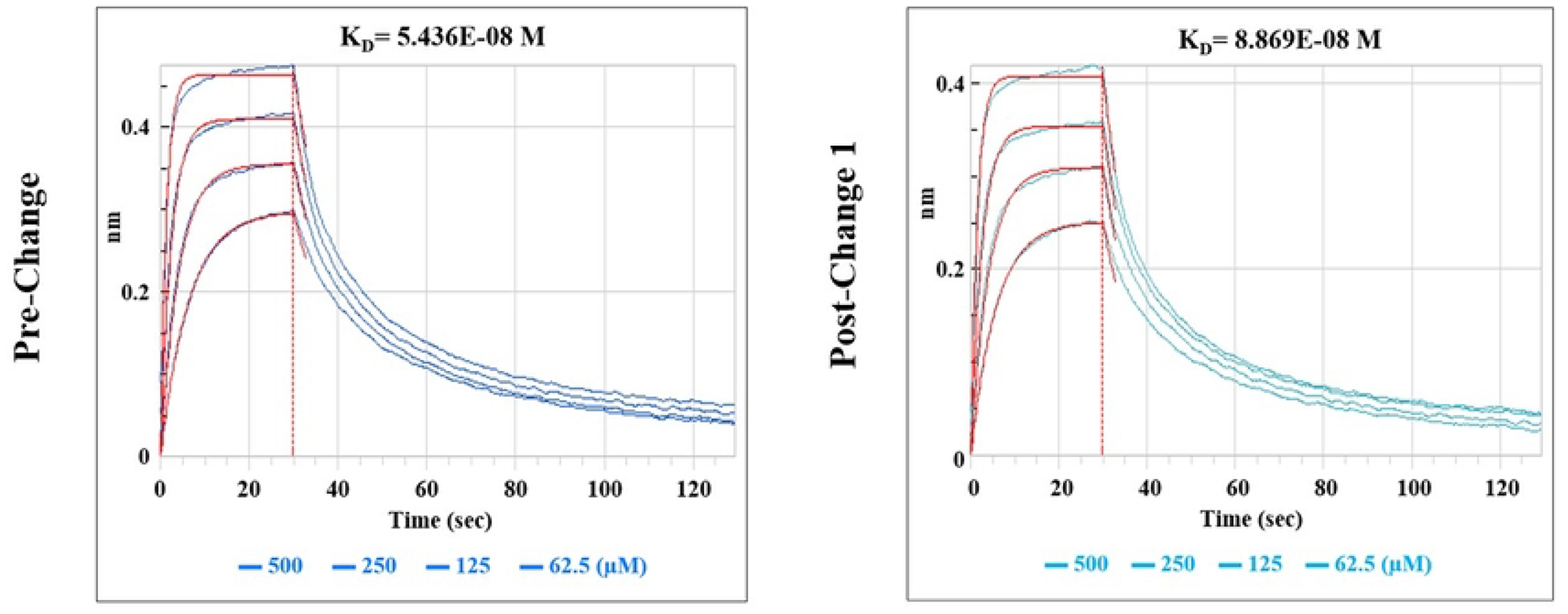
Side-by-side effector function characterization of ARX788 DS batches: A) ADCC Dose Response Curves, B) Fc gamma Receptors Binding Kinetics Sensorgrams, and C) FcRn Receptors Binding Kinetics Sensorgram

#### Side-by-Side Forced Degradation Study

To ascertain comparative stability and comparability under thermal stress, DP lots were incubated at an elevated temperature of 40°C for 1, 2, and 4 weeks in an upright position of the vials. Prior studies of forced degradation including freeze/thaw, agitation, and thermal stress had indicated only thermal stress as the most aggressive stress condition. Therefore, only thermal stress condition was used in the comparability study. Since the pre-change DP is in liquid form while post-change DP is lyophilized powder, a set of post-change1-3 DP vials were reconstituted at T0 to ∼10 mg/mL (same concentration as pre-change DP) and then incubate as liquid DP together with pre-change DP at 40°C with 75±5% relative humidity, to ensure a fair comparison for the forced degradation study. Another set of post-change1-3 DP vials were incubated under the same thermal stress condition without reconstitution to determine if the lyophilized dosage form would improve ARX788 DP stability. The degradation profiles of ARX788 DP lots were then determined by a side-by-side comparison of key quality attributes using stability indicating batch release analytical and biological assays (latter not shown). In addition, the T0 data in the forced degradation study allowed assessment of the side-by-side batch release testing requirement in ICH Q5E guidance. Only representative data from the Pre-change vs. Post-change Lyo 1 batch are shown.

##### CEX-HPLC

The level of charge heterogeneity in both lots of thermally stressed drug product from pre-change and post-change was determined by CEX-HPLC. The magnified CEX chromatograms of pre-change, reconstituted post-change Lyo 1 and post-change Lyo 1 are shown in Fig5A. A clear trend in reduction of main peak purity and concomitant appearance of acidic and basic species for both pre-change and reconstituted post-change Lyo 1 can be observed, while the lyophilized post-change DP purity remains unchanged. From the chromatograms, it can also be observed that the rates of change were very comparable. Overall, this indicates that the drug product lots from both Pre-change and Post-change Lyo (after reconstitution) degrade similarly when thermally stressed at 40°C and signifies a high degree of similarity of the ADC between these two Processes. Post-change Lyo DP in its lyophilized state is resistant to this degradation, which shows the improved stability of this new dosage form.

**Figure 5:**
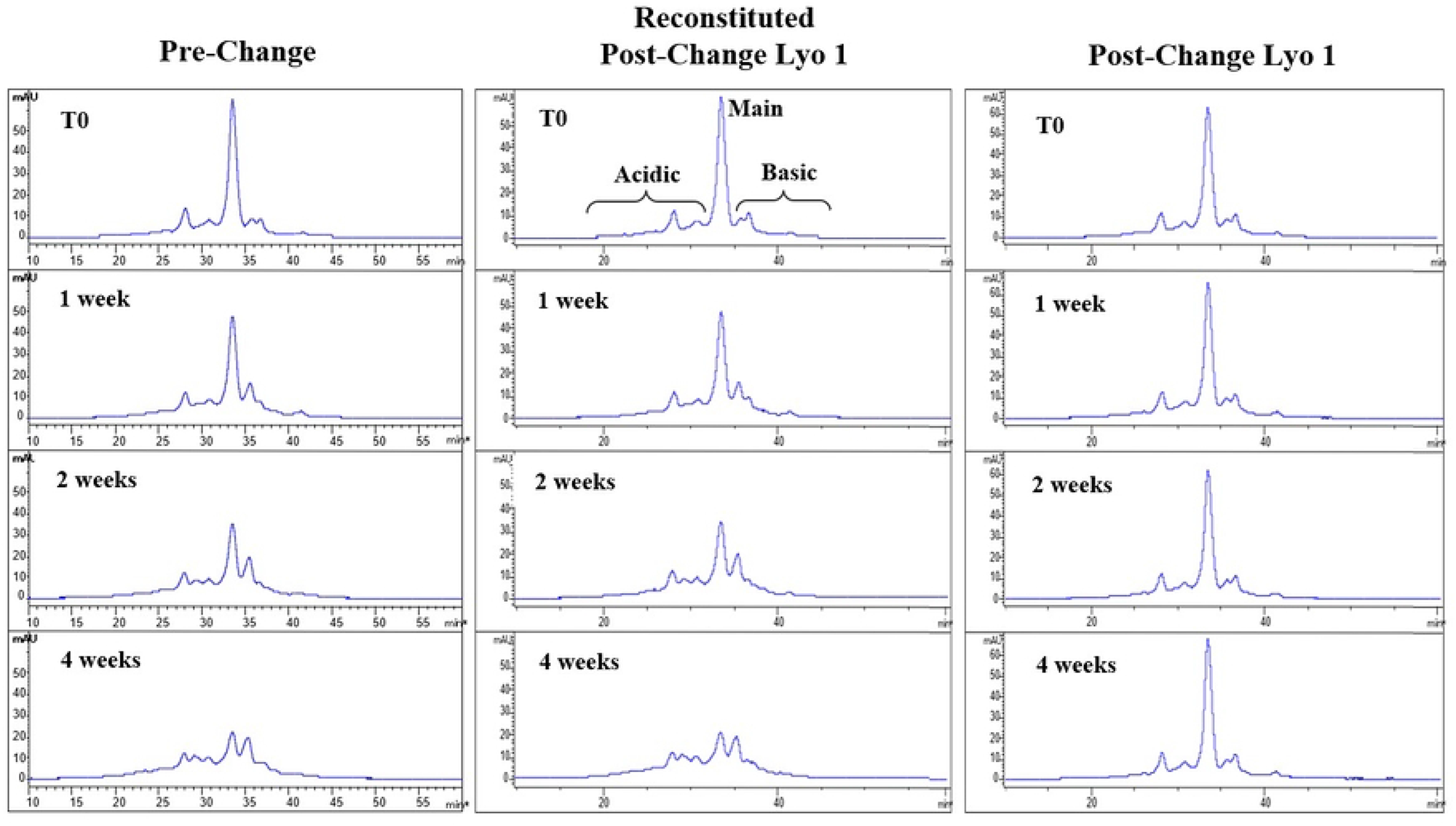
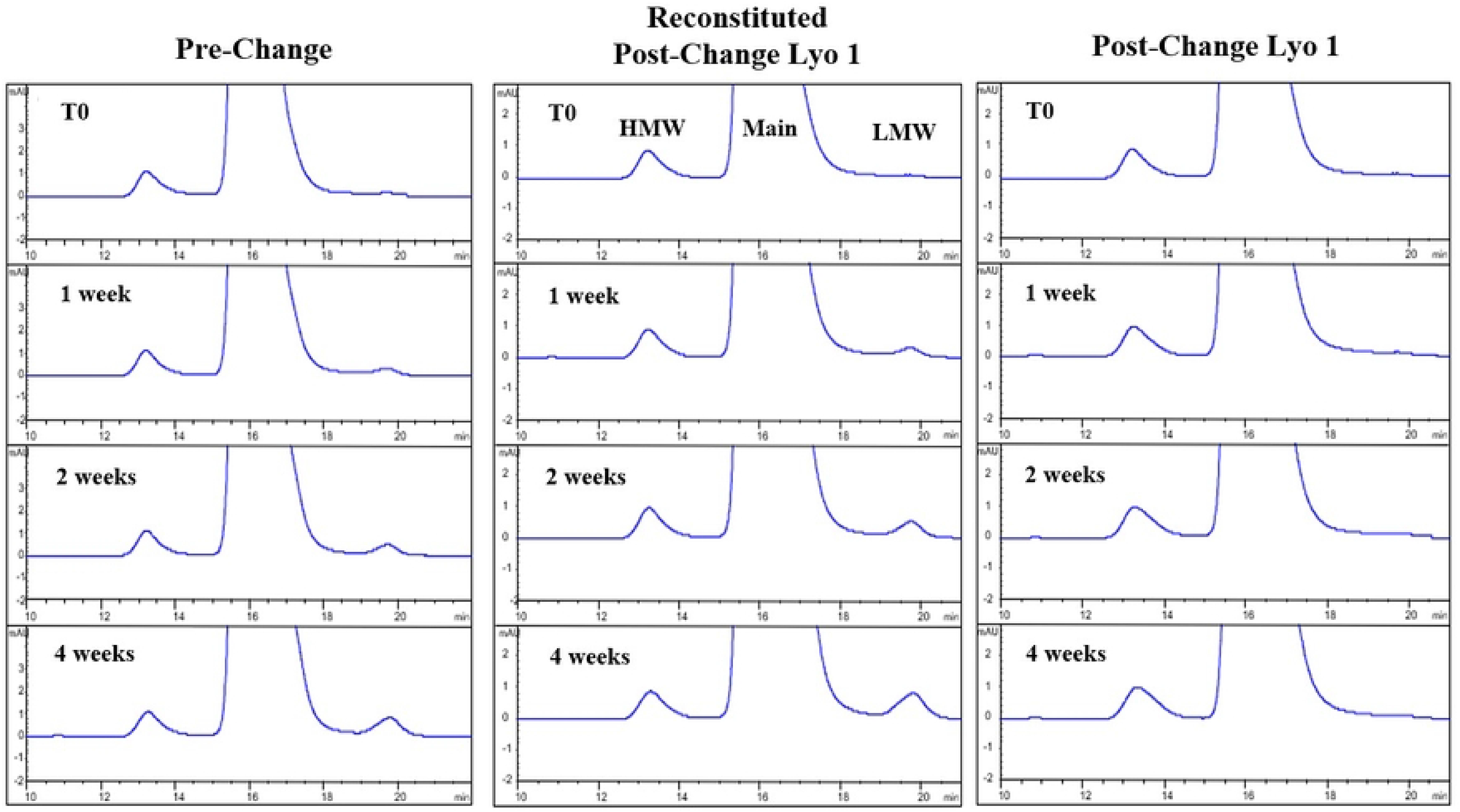
Side-by-side forced degradation purity analysis of ARX788 DP lots by A) CEX HPLC, B) SEC HPLC (Green arrow indicates LMW peak)

##### SEC-HPLC

The level of HMW, monomeric, and LMW species of the thermally stressed drug product was determined by SE-HPLC. The magnified SEC chromatograms of control samples (T0) and all three timepoints are shown in Fig 5B. Pre-change and reconstituted Post-change samples both showed a similar gradual increase in LMW. Drug product from both Pre-change and Post-change Lyo show identical trend with gradual drop in main peak purity and increase in LMW levels indicating high degree of comparability between these two processes. Post-change in its lyophilized state is resistant to this degradation, which shows the improved stability of this new dosage form.

##### HIC-HPLC

The drug to antibody ratio (DAR), along with the unconjugated mAb, 1-drug, 2-drug, and impurities in the thermally stressed (up to 4 weeks at 40°C) drug product lots were determined by HIC-HPLC. The data are shown in Table 10. The results show that after 4 weeks of thermal stress, Pre-change and reconstituted Post-change 1 Lyo drug product lots show a small increase in relative abundance of DAR1 species and a minute decrease in the relative abundance of DAR2 species indicating gradual loss of the AS269 drug. These results correlate well with free drug assay results where a minute increase in free drug was observed under identical forced degradation conditions, namely 4 weeks at 40°C (see below). The nominal DAR for all lots of thermally stressed DP however remains the same (1.9), as the changes were very small. The high degree of similarity in the HIC-HPLC results indicate that the drug-antibody conjugate profile and stability of the Pre-change and Post-change lot after reconstitution, are comparable. The non-reconstituted Post-change lots are resistant to this degradation and the free drug assay also shows that there is no increase in free drug levels (see below). These data also show that the oxime bond that conjugates the drug linker payload, AS269, to the antibody is very stable under heat stress in the lyophilized DP.

**Table 10:** Calculated DAR of Thermally Stressed ARX788 Drug Product by HIC-HPLC

| Sample | species | T0 | Week 1 | Week 2 | Week 4 |
| --- | --- | --- | --- | --- | --- |
| Pre-Change | % mAb | 0.2 | 0.2 | 0.3 | 0.3 |
|  | % 1-Drug | 8.1 | 8.9 | 9.1 | 10.7 |
|  | % 2-Drugs | 85.4 | 85.4 | 85.6 | 83.8 |
|  | % Impurities | 6.2 | 5.6 | 5.0 | 5.2 |
|  | DAR | 1.9 | 1.9 | 1.9 | 1.9 |
| Post-Change 1 -<br>Reconstituted | % mAb | 0.3 | 0.3 | 0.4 | 0.5 |
|  | % 1-Drug | 11.0 | 11.6 | 11.7 | 13.1 |
|  | % 2-Drugs | 83.0 | 83.3 | 83.1 | 81.6 |
|  | % Impurities | 5.7 | 4.8 | 4.7 | 4.8 |
|  | DAR | 1.9 | 1.9 | 1.9 | 1.9 |
| Post-Change 1 -<br>Lyophilized DP | % mAb | 0.3 | 0.3 | 0.5 | 0.4 |
|  | % 1-Drug | 11.0 | 11.0 | 10.6 | 10.8 |
|  | % 2-Drugs | 83.0 | 83.6 | 83.6 | 84.0 |
|  | % Impurities | 5.7 | 5.1 | 5.4 | 4.8 |
|  | DAR | 1.9 | 1.9 | 1.9 | 1.9 |

### Residual Free Drug analysis

The level of free drug impurity in thermally stressed ARX788 drug product lots was determined by RP-HPLC assay as described in the Materials and Methods section. The amount of free drug (shown as calculated % weight of free drug per weight of ADC) for Pre-change, Post-Change 1 DP reconstituted, and Post-Change 1 Lyophylized DP are listed in table 11. The data show that both Pre-Change and reconstituted Post-Change 1 DP demonstrate a similar slow rate of increase in level of free drug. This indicates that when subjected to thermal stress at 40°C, the pre-change and post change DP lots degrade in a similar manner, demonstrating a high degree of comparability. Post-Change DP in the lyophilized state before reconstitution is resistant to this degradation, which demonstrates the improved stability of this new dosage form.

**Table 11.** Free Drug Impurity in Thermally Stressed ARX788 Drug Product.

| Sample | T0 | Week 1 | Week 2 | Week 4 |
| --- | --- | --- | --- | --- |
| Pre-Change | <0.016 | <0.016 | 0.023 | 0.032 |
| Post-Change 1 - Reconstituted | <0.016 | 0.020 | 0.026 | 0.037 |
| Post-Change 1 -Lyophilized DP | <0.016 | <0.016 | <0.016 | <0.016 |

Overall, the forced degradation profiles assessed using stability indicating analytical assays and biological activity assays (data not shown) of DS and DP from pre-change and post-change batches at 40°C were highly similar.

## Conclusion

In summary, side-by-side comparison of pre-change and post-change DS and DP was conducted by evaluating different chemical, biochemical and biological functions and included a comparison of degradation profiles under thermal stress conditions. All results have demonstrated a high degree of similarity between the pre- and post-change ARX788 materials. Further, as part of the product quality comparability assessment, results from long term and accelerated stability studies evaluated to date have shown high similarity in purity and impurity profiles (data not shown). Therefore, it is anticipated that the changes made to optimize the ARX788 manufacturing process will not adversely impact product quality, safety, pharmacokinetics, potency, and efficacy.

## Acknowledgments

The Authors thank Ji Young Kim for her supervision of the ADCC effector function analysis, and Karen Cha for her expert guidance on global regulatory submissions.

